# Tyrosine-sulfated peptide hormone induces flavonol biosynthesis to control elongation and differentiation in Arabidopsis primary root

**DOI:** 10.1101/2024.02.02.578681

**Authors:** Maria Florencia Ercoli, Alexandra M. Shigenaga, Artur Teixeira de Araujo, Rashmi Jain, Pamela C. Ronald

## Abstract

In Arabidopsis roots, growth initiation and cessation are organized into distinct zones. How regulatory mechanisms are integrated to coordinate these processes and maintain proper growth progression over time is not well understood. Here, we demonstrate that the peptide hormone PLANT PEPTIDE CONTAINING SULFATED TYROSINE 1 (PSY1) promotes root growth by controlling cell elongation. Higher levels of PSY1 lead to longer differentiated cells with a shootward displacement of characteristics common to mature cells. PSY1 activates genes involved in the biosynthesis of flavonols, a group of plant-specific secondary metabolites. Using genetic and chemical approaches, we show that flavonols are required for PSY1 function. Flavonol accumulation downstream of PSY1 occurs in the differentiation zone, where PSY1 also reduces auxin and reactive oxygen species (ROS) activity. These findings support a model where PSY1 signals the developmental-specific accumulation of secondary metabolites to regulate the extent of cell elongation and the overall progression to maturation.

Teaser

PSY1-induced flavonol biosynthesis in Arabidopsis roots modulates the distance from the root tip at which cell elongation ceases.

## Introduction

In multicellular organisms, growth involves cell proliferation and expansion. The shape and final dimensions of an organ are determined by the balance between growth initiation and cessation. In roots, these dynamics underlie primary growth, causing the root to extend along its longitudinal axis (*1*). Although it is known that the establishment and maintenance of developmental boundaries are critical for the spatiotemporal regulation of growth initiation and cessation (*2*), how different regulatory networks are integrated to control the magnitude of cellular growth in roots is not well understood.

In the *Arabidopsis thaliana* primary root, cellular growth is initiated, maintained, and eventually terminated (Fig. 1A, left side of the panel) (*3*). The different cell types that constitute the root arise from generative cell divisions of stem cells located in the stem cell niche (SCN) located at the proximal (rootward) end of the root tip. The SCN is maintained by a small group of slowly dividing organizer cells known as the quiescent center (QC) (*4*). More distally (shootward) from the QC, in the meristematic zone (MZ), proliferating cells provide the necessary number of cells for organ growth . Cells stop dividing at the transition zone (TZ) but continue to grow rapidly by directional expansion in the adjacent elongation zone (EZ) (*6–8*). Cell elongation slows down in the distal parts of the EZ in what is defined as the start of growth cessation (*9*). Finally, growth ceases in the differentiation zone (DZ), and cells mature into their final shape and function (*10*). Because postembryonic root growth is indeterminate, these processes are continual, resulting in a developmental gradient along the root longitudinal axis, referred to as root zonation (Fig. 1A, left side of the panel) (*11*).

**Figure 1.**
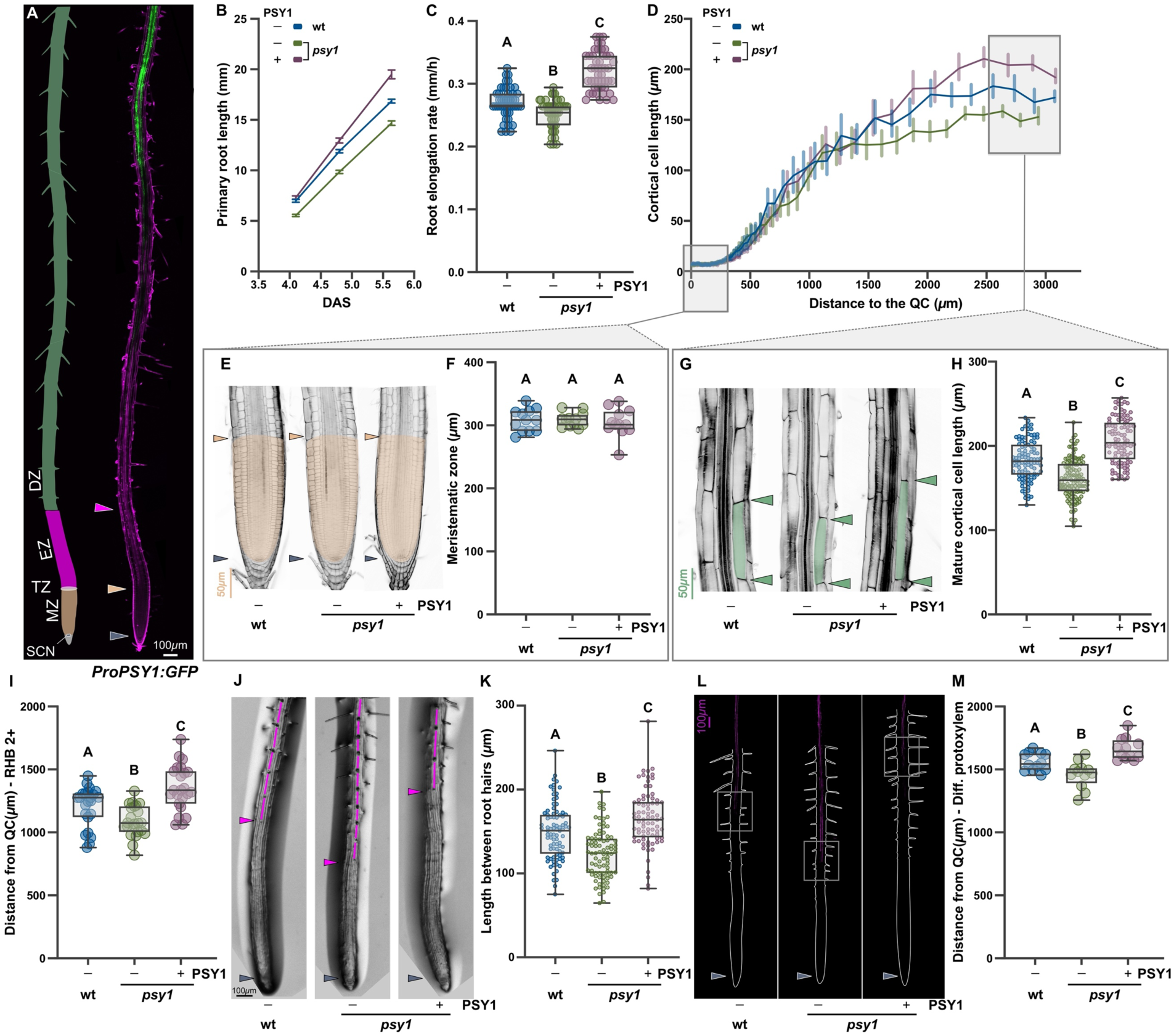
Loss of PSY1 impairs Arabidopsis root growth. (**A**) A scheme describing *Arabidopsis thaliana* root zonation (left side of the panel). SCN, stem cell niche; MZ, meristematic zone; EZ, elongation zone; and DZ, differentiation zone. A 7-day-old Arabidopsis primary root expressing a *PSY1* promoter-GFP transcriptional reporter (*ProPSY1:GFP*) (green). Cell walls were stained with propidium iodide (PI, magenta). (**B**) Root growth, (**C**) root elongation rate (mm/h)(**n=50** seedlings), and (**D**) cortical cell length profile (**n=12** seedlings) in wild-type (wt) and *psy1* seedlings grown for 6 days on 1xMS vertical plates with or without 50nM of synthetic PSY1. The meristematic zone size (**E** and **F, n=12** seedlings) and mature cortical cell length (**G** and **H, n=100** cells) are highlighted. The limits of a representative meristematic zone (**E**) and mature cortical cells (**G**) are shaded in pale orange and green, respectively. (**I**) Distance from QC to the first root hair bulge at stage +2 (RHB 2+) (**n=23** seedlings), (**J**) root tip architecture, (**K**) length between consecutive root hairs in one trichoblast file (**n=80**), (**L** and **M, n=12** seedlings) distance from QC to differentiated vascular elements (Diff. protoxylem) revealed by basic fuchsin staining from *wt* and *psy1* seedlings grown for 6 days on 1xMS vertical plates with or without 50nM of synthetic PSY1. In (**J**), the length between consecutive root hairs in one trichoblast file is highlighted in magenta. In (**L**), the grey squares indicate the zone where the deposition of lignin starts in the protoxylem, as revealed with basic fuchsin staining, and organ boundaries are marked by white dashed lines. In (**C**), (**F**), (**H**), (**I**), (**K**), and (**M**), the data shown are box and whisker plots combined with scatter plots; each dot indicates the measurement of the designated parameter listed on the y-axis of the plot. Different letters indicate significant differences, as determined by one-way ANOVA followed by Tukey’s multiple comparison test (P < 0.05). The purple arrowheads mark the position of the QC, the pale orange arrowheads mark the end of the meristem, where cells start to elongate, the green arrowheads indicate the mature cortical cell size, and the magenta arrowhead points to the first root hair bulge, defined as stage +2, indicating the end of the elongation zone in the epidermis.

To ensure proper root zonation, a combination of mechanical forces, transcriptional regulators, phytohormone, and metabolic inputs interact to establish the developmental boundaries that maintain the balance between growth initiation and cessation (*12*). The transition between the MZ and EZ involves a complex interplay of phytohormones, primarily auxin and cytokinin (*8*). Polar auxin transport (PAT) generates a gradient in the root with a maximum at the SCN. This gradient regulates the distribution of *PLETHORA* (*PLT*) transcription factors (*13*, *14*). PLT proteins display a graded distribution and modulate root zonation in a dosage-dependent manner: high levels are required for maintenance of the SCN, intermediate levels induce rapid cell divisions in the MZ, and low levels facilitate cell elongation and differentiation (*15–17*). Cytokinin controls PAT and auxin degradation, generating an auxin minimum precisely positioning the TZ (*7*). In the elongation/differentiation transition, distinct mature cellular characteristics such as cell wall structure (*18*), microtubule orientation (*19*), root hair development in the epidermis (*20*), and lignified secondary cell walls in the protoxylem (*21*), among others, point to a pronounced shift in cell identity and function across these root regions. The interplay between regulatory mechanisms underpinning this second developmental boundary where the cessation of rapid cell elongation and maturation of root cells occurs remains underexplored.

Alongside classic plant hormones, small tyrosine-sulfated peptides play significant roles in the complex regulatory networks controlling root growth and are thus candidates for signaling in the developmental trajectories in root zonation (*22*, *23*). Functioning as extracellular signals, these peptide hormones synchronize cellular activities across tissues. Typically, they bind to leucine-rich repeat receptor-like kinases (LRR-RLKs) to trigger specific signaling pathways (*24*). Sulfation of a tyrosine residue in the peptide sequence, a posttranslational modification carried out by *TYROSYLPROTEIN SULFOTRANSFERASE* (*TPST*), is required for signaling activity, as it regulates peptide affinity for its cognate receptor (*25–28*). An example is the nine-member *PLANT PEPTIDE CONTAINING SULFATED TYROSINE* (*PSY*)-family in Arabidopsis (*29*, *30*). To exert their function, these peptides bind to three cognate LRR-RLKs known as *PSYR* or *ROOT ELONGATION RECEPTOR KINASES (REKs)1/2/3* (*30*, *31*). The triple PSYR knockout (*psyr123 or tri-1*) displays an elongated root phenotype compared with wild-type plants, suggesting that PSYR1/2/3 signaling negatively regulates root growth (*30*, *31*). The application of synthetic PSY peptides enhances root growth of wild-type and *tpst* knockout plants but not the triple receptor mutant, supporting a role for PSYR1/2/3 as receptors for PSY peptides (*29*, *30*, *32*). Among this peptide family, PSY1 is the most extensively studied, particularly its association with the regulation of mature cell size in the root cortex and seedling cuticle development (*29*, *33*). Despite these advances, the broader significance of PSY signaling in plant root growth remains to be fully characterized.

In this study, we demonstrate that PSY1 regulates root growth, controlling the magnitude that cells elongate before reaching their final, differentiated size. Transcriptomic analysis conducted on Arabidopsis roots treated with synthetic PSY1 revealed an upregulation of genes controlling the biosynthesis of flavonols, a class of plant-specific secondary metabolites. Flavonol-specific staining and analysis of the expression pattern of flavonoid biosynthetic enzymes indicate that these metabolites accumulate in the DZ upon PSY1 treatment. Genetic and chemical treatments provide evidence that flavonol biosynthesis is required for PSY1-induced root growth. Finally, we found that auxin activity and ROS accumulation vary according to the PSY1 abundance along the longitudinal axis of the root, suggesting a role for PSY1 in controlling the distance from the QC at which cell elongation slows down in different root tissues. Together, our findings demonstrate that root zonation requires spatial regulation of flavonol accumulation through a developmental-specific expression of genes encoding biosynthetic enzymes. These results significantly advance our understanding of the mechanisms that control cell elongation and differentiation in Arabidopsis roots.

## Results

### 1. PSY1 controls cell elongation and differentiation in primary roots

To explore the role of PSY1 in Arabidopsis primary root growth, we examined PSY1 promoter expression in Arabidopsis roots. We found that PSY1 is highly expressed in the DZ as determined using publicly available gene expression profiles of manually dissected root tissue segments corresponding to MZ, EZ, and DZ (*34*)(fig.S1A). A matching expression profile was obtained when utilizing the single-cell Arabidopsis root atlas (*35*)(fig.S1B). To validate these results, we generated 15 independent transgenic lines expressing the transcriptional reporter *ProPSY1:GFP* (PSY1 promoter-driven GFP) in the wild-type Col-0 (wt) background. We observed that PSY1 promoter activity gradually increased in the progression of the DZ, with the GFP signal starting to rise approximately 2000μm from the QC (Fig.1A, and figS1.C). Interestingly, when we analyzed *ProPSY1:GFP* expression in the DZ, we found that the GFP signal was almost undetectable in the epidermis, consistent with the pattern of PSY1 expression in various root tissues at different developmental stages, as documented in the single-cell Arabidopsis root atlas (*35*)(fig.S1, B and D).

To investigate the function of PSY1 in root development, we generated ectopic expression lines in a wt background using the constitutive 35S promoter (*Pro35S:PSY1*). Consistent with previous reports, these plants developed longer primary roots (fig.S2, A-C) (*29*). The same phenotype was observed when wt plants were treated with synthetic PSY1 (fig.S3, A-C). Exogenous application of PSY1 can also partially restore root growth in the tyrosylprotein sulfotransferase mutant, *tpst-1*, which is deficient in biosynthesis of all tyrosine sulfated peptides (fig.S3, A-C) (*25*, *36*). Conversely, the *psy1* knockout (*33*) displayed reduced root length and elongation rates (Fig.1, B and C). Synthetic PSY1 treatment rescued the root growth defect in the *psy1*, resulting in a phenotype resembling that of wt roots subject to PSY1 treatment (Fig.1, B and C, and fig.S3, A-C).

To further examine the root growth phenotype, we constructed a cell length profile by measuring the length of individual cortical cells from the QC to the DZ. In a typical cell length profile, cell length remains relatively short and constant in the MZ, sharply increases in the EZ, and eventually levels off in the DZ, where cells attain their final size and identity (*3*). Our profile analysis revealed that the MZ length in *psy1* and *Pro35S:PSY1* are indistinguishable from wt plants (Fig.1, D-F, and fig. S2, D-F). Additionally, the short MZ in *tpst-1* mutants remained unaffected when grown in media supplemented with synthetic PSY1 (fig.S3, D-F). Consistent with these findings, the expression and distribution of the G2-to-mitosis transition marker *CYCLINB1;1*, commonly used to assess cell proliferation in the MZ (*3*), were unchanged in the plants expressing ectopic PSY1 (fig.S4, A-D). These results indicate that PSY1 does not regulate cell proliferation in the MZ.

Because the establishment of the TZ relies on the auxin/*PLETHORA*/cytokinin regulatory node (*8*, *16*, *37*), we investigated the responses of this molecular network following synthetic PSY1 treatment. We observed no significant differences in the intensity and distribution of auxin and cytokinin response reporter lines (*DR5v2:n3GFP* (*38*) and *pTCSn::GFP* (*39*), respectively) compared with untreated plants (fig.S4, E-H). Additionally, there were no differences in localization and expression of *PLETHORA1* (*PLT1*) between PSY1-treated and untreated plants based on our analysis of a transcriptional reporter line (*ProPLT1:CFP*) (fig.S4, I-K). Because *PLT1* expression is also controlled post-translationally (*36*, *40*), we evaluated the response of a plant expressing translational fusion (*ProPLT1-PLT1:YFP)*. *ProPLT1-PLT1:YFP* expression did not change in response to PSY1 treatment (fig.S4, L-N). In contrast, when treated with ROOT GROWTH FACTOR 1 (RGF1), a small tyrosine-sulfated peptide known to regulate MZ size by stabilizing PLT proteins, the PLT1-YFP signal showed an enhanced and broader expression in the MZ. Together, these results show that PSY1 does not modify the position of the TZ and, therefore, does not affect the size of the MZ.

We next leveraged the cell length profiles to investigate the role of PSY1 in the control of cell elongation. We found that where the cortical cells in the *psy1* mutant reached their final cell size, cortical cells were still elongating in the wt plant, suggesting a premature exit from elongation in the mutant (Fig.1D). Consistent with this, *psy1* mutants exhibited significantly shorter mature cortical cells compared to wt (Fig.1, D, G, and H). Conversely, when *psy1* mutants were grown in media supplemented with synthetic PSY1, the cortical cells continued to elongate, whereas cells in the wt stalled, resulting in significantly longer mature cortical cell sizes compared to those of wt plants (Fig.1, D, G, and H). Similar effects were observed in *tpst-1* plants grown in media supplemented with PSY1, as well as in wt plants expressing *Pro35S:PSY1* (fig.S2, D, G, and H, and fig.S3, D, G, and H) These findings denote a role for PSY1 primarily controlling root growth by defining the extent to which cells elongate before cells reach their final, differentiated size.

If our hypothesis is correct, PSY1 may also affect the distance from the QC at which cell elongation slows down in other tissues. In the trichoblast cell files of the epidermis, the onset of root hair development marks the cessation of rapid cell elongation, followed by minimal further cell elongation as these cells attain their mature size (*41*). To explore this in the context of PSY1 signaling, we measured the distance from the root tip to where the first root hair bulge at stage +2 can be identified (*20*). Notably, in *psy1* mutants, we observed that this distance is reduced compared with wt plants (Fig.1, I and J). In line with a premature appearance of root hairs and exit of cell elongation, the length between consecutive root hairs in one trichoblast file was reduced in *psy1* (Fig.1, J and K). Additionally, secondary cell wall formation, as evidenced by the characteristic helical lignin pattern, a maturation cue in the tracheary elements of protoxylem (PX)(*42*), appeared closer to the root tip of *psy1* plants as visualized using basic fuchsin staining (Fig.1, L and M). These findings suggested that in *psy1* mutants, hallmarks of maturation in different tissues have shifted toward the root tip. In contrast, synthetic PSY1 treatment of *psy1*, wt, and *tpst-1* plants or PSY1 overexpression led to root hair initiation farther from the root tip, increased distance between root hairs in a single trichoblast cell file, and shootward displacement of lignin deposition in PX (Fig.1, I-M; fig.S2, I-M and fig.S3, I-M). Altogether, these results indicate that PSY1 functions as a crucial signal necessary for a normal root zonation.

### 2. PSY1 regulates genes highly expressed in elongation and differentiation zones

To investigate how PSY1 controls root zonation, we carried out RNA-sequencing (RNA-seq) profiling of roots treated with synthetic PSY1. For these experiments, we harvested the root tips, including the DZ, from 5-day-old wt seedlings treated with PSY1 for 4 hours (Fig.2A). Because seedlings need to be treated with synthetic PSY1 for at least 24 hours to exhibit root zonation changes (fig.S5, A and B), we can rule out the possibility that these anatomical changes are the cause of any RNA level modification in this experiment. The RNA-seq analysis revealed that 253 genes (92 activated, 161 repressed) were differentially expressed after 4 hours of PSY1 treatment (Supplemental Data Set 1).

**Figure 2.**
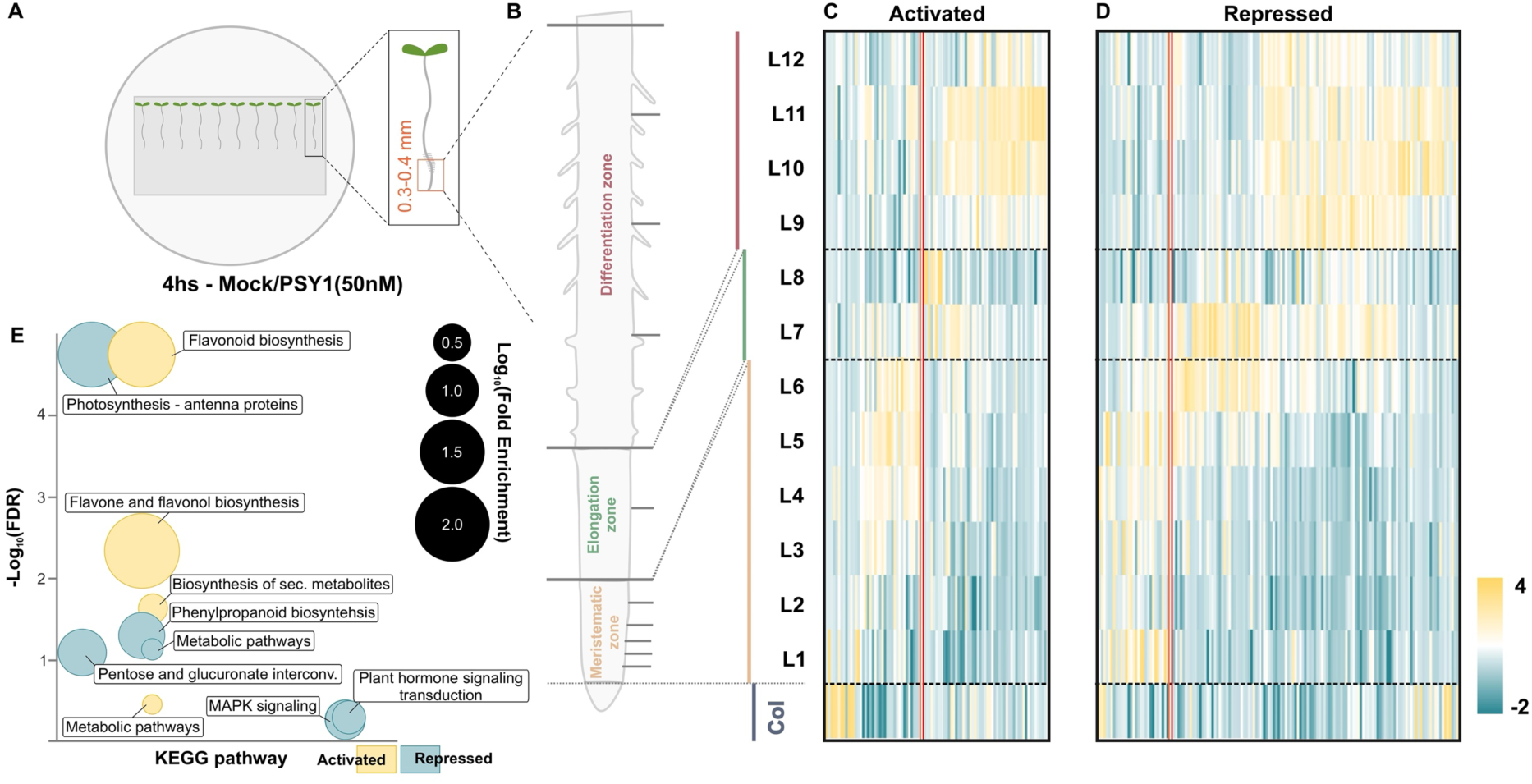
PSY1 regulates genes highly expressed in the elongation and differentiation zones. (**A**) Seedlings were grown on a permeable mesh placed on a clear agar substrate. After 5-d, seedlings were transferred to a control medium or medium containing PSY1 (50nM) by moving the mesh to the selected media. After 4 hours, samples were harvested. RNA was extracted from designated root zones. (**B**) Scheme of a root showing developmental zones. Horizontal lines define the sections (L1-L12) of the longitudinal Root Gene Expression Atlas (Brady et al., 2007). Col, Columella. Sections 1 to 6 comprise the Meristematic zone; 7 and 8, the Elongation zone; and 9 to 12, the Differentiation zone. Expression along the root’s longitudinal axis of genes activated (**C**) or repressed (**D**) after PSY1 treatment. Z-scores were calculated across samples and expressed as a heat map. The double red line separates genes preferentially expressed in columella and meristematic regions (left) from those that present a maximum expression in the elongation and differentiation zone (right). (**E**) Bubble plot for Kyoto Encyclopedia of Genes and Genomes (KEGG) pathway enrichment analysis of genes activated or repressed after PSY1 treatment <u>(Supplemental Data Set 3</u>). The y-axis shows the False Discovery Rate (FDR) in a negative Log10 scale, whereas the x-axis is fixed, and terms from the same KEGG subtree are located closer to each other. The size of each circle represents the term Fold Enrichment in the Log10 scale.

Given the role of PSY1 controlling growth by determining the magnitude of cell elongation and mature cell size without affecting cell proliferation, we anticipated that genes downstream of PSY1 signaling would primarily be expressed outside the MZ. To test this hypothesis, the expression patterns of the PSY1-responsive genes were analyzed in the developmental stage-specific gene expression database (*34*) (Fig.2, B-D). We found that 57% of the PSY1-activated genes (51 out of 89 genes) (Fig.2C) and 80% of the PSY1-repressed genes (117 of 146 genes) (Fig. 2D) exhibited a peak of expression in a region corresponding to the EZ and DZ. This spatial pattern shows that PSY1 controls genes preferentially expressed in the same zones of the root where the PSY1-associated phenotypes are observed.

Gene Ontology (GO) and KEGG pathway enrichment analyses were conducted to explore the functional implications of PSY1-responsive genes (Fig.2E, fig.S5, C and D and Supplemental Data Set 2 and 3). Notably, secondary metabolic processes and flavonol biosynthetic pathways were significantly enriched categories among the PSY1-activated genes (Fig.2E, fig.S5C and Supplemental Data Set 2 and 3). Flavonols, a class of flavonoids that are secondary metabolites in plants, include compounds such as quercetin and kaempferol, along with their glycosylated derivatives, which are commonly found in Arabidopsis roots (*43*). Enzymes responsible for synthesizing these two specific flavonol scaffolds were upregulated in response to PSY1 treatment (Fig.3 and Supplemental Data Set 1). Additionally, the transcription factor *MYB12*, which is known to control flavonol biosynthesis primarily in the root (*44*, *45*), the multidrug and toxin efflux flavonoid transporter *DTX35* (*46*), the *RHAMNOSE SYNTHASE RHM1/ROL1* (*47*), and the *3-KETOACYL-COA THIOLASE* isoform *KAT5* (*48*) were also upregulated by PSY1 treatment (Fig.3 and Supplemental Data Set 1). *KAT5* and *RHM1* were previously shown to be closely associated with genes involved in flavonoid biosynthesis in a co-expression analysis (*49*). Furthermore, our time course study of whole seedlings treated with synthetic PSY1 coupled with quantitative reverse transcription polymerase chain reaction (RT-qPCR) demonstrated that a subset of genes differentially regulated after 4 hours of PSY1 treatment maintained their expression levels at 8, 12, and 48 hours (fig.S5G).

**Figure 3.**
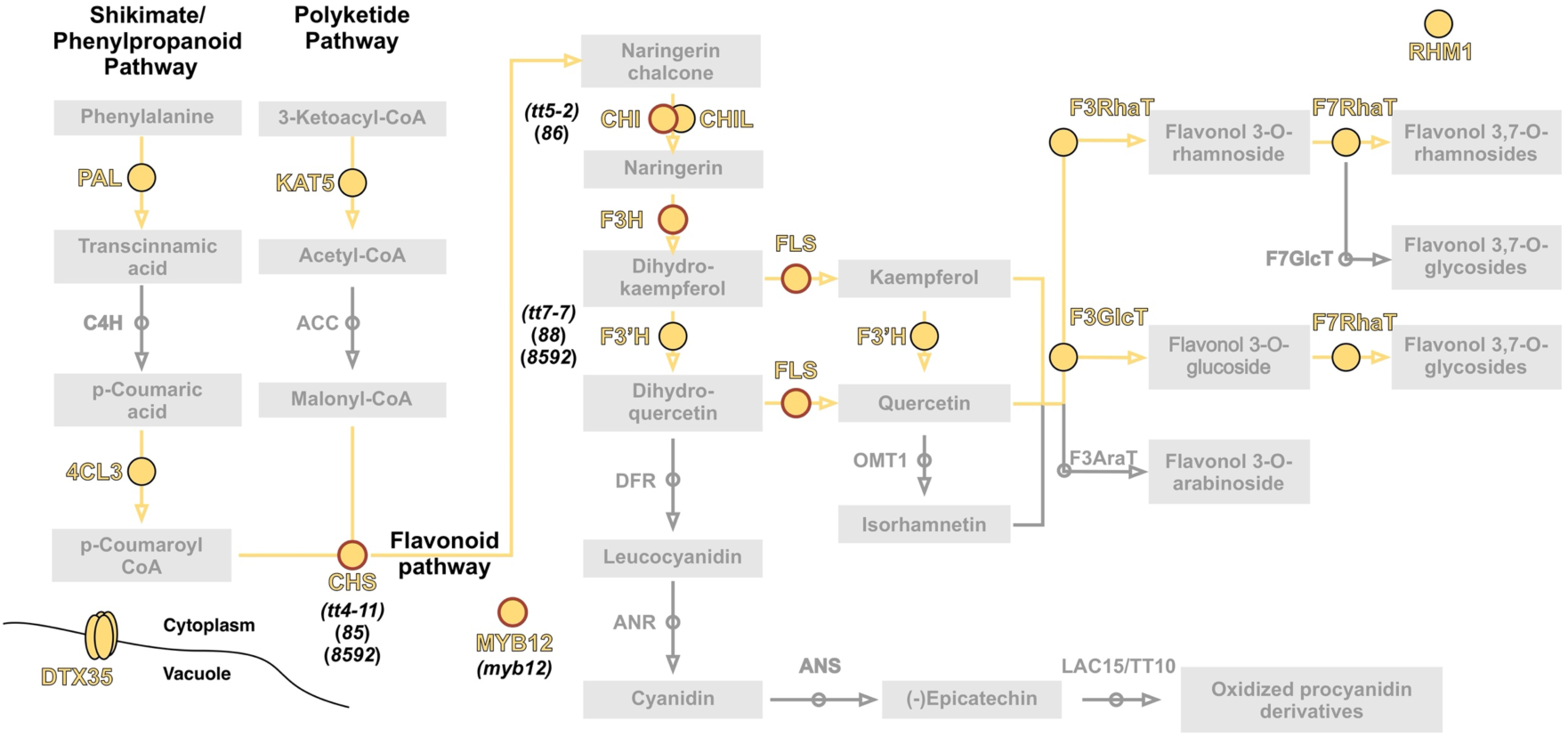
PSY1 regulates genes that code for enzymes involved in the production of flavonols. Flavonoid biosynthetic pathway. Circles represent the flavonoid biosynthesis enzymes. The metabolites are labeled with rectangular boxes. Highlighted in larger yellow circles are the genes coding for enzymes activated after PSY1 treatment. Yellow circles with red borders denote the genes coding for enzymes that are also positively regulated by MYB12, according to (45). *PAL,* phenylalanine ammonia-lyase; *C4H,* cinnamic acid 4-hydroxylase; *4CL,* 4-coumaric acid: CoA ligase; *KAT5,* 3-ketoacyl-coa thiolase; *ACC,* acetyl-CoA carboxylase; *CHS,* chalcone synthase; *CHI/CHIL,* chalcone isomerase*; F3H,* flavanone 3-hydroxylase*; F3’H,* flavonoid 3-hydroxylase*; FLS,* flavonol synthase*; OMT1,* o-methyltransferase 1*; DFR,* dihydroflavonol 4-reductase*; ANS,* anthocyanidin synthase*; ANR,* anthocyanidin reductase*. F3ARAT,* flavonol 3-o-arabinosyltransferase*; F3GLCT,* flavonoid 3-o-glucosyltransferase*; F3RHAT,* flavonol 3-o-rhamnosyltransferase*; F7GLCT,* flavonol 7-o-glucosyltransferase*; F7RHAT,* flavonol 7-o-rhamnosyltransferase*; RHM1,* UDP-rhamnose synthase 1*; DTX35,* detoxifying efflux carrier 35*; MYB12, R2R3-MYB*. Names of the mutant lines of genes coding for enzymes used in subsequent experiments are in parentheses (<u>Supplemental Table 1</u>).

Recent research has suggested that PSY-family peptides play a role in repressing PSYR function, thereby facilitating growth (*30*). To explore this further, we investigated whether synthetic PSY1 treatment could mimic the effects of PSYR loss at the molecular level. We conducted KEGG pathway enrichment analysis using the 1,947 genes activated in the triple PSYRs mutant background, *tri-1,* as reported by (*31*). Notably, categories related to flavonoid and phenylpropanoid biosynthesis were significantly enriched (fig.S5E and Supplemental Data Set 4). Additionally, among the 41 genes activated following both PSY1 treatment and the triple PSYR mutant, 13 were associated with flavonol biosynthesis (fig. S5F). Collectively, these findings indicate that synthetic PSY1 treatment and PSYR loss both activate genes involved in the biosynthesis of secondary metabolite intermediates.

### 3. PSY1 regulates flavonol accumulation in the differentiation zone

Previous studies have revealed that flavonoid accumulation is developmentally regulated and occurs in a tissue-specific manner, matching the expression pattern of genes involved in early flavonoid biosynthesis (*50*, *51*). Using publicly available transcriptomics data sets of manually dissected root tissue segments corresponding to MZ, EZ, and DZ (*34*), we found that the expression of genes that produce the majority of flavonols, including *CHALCONE SYNTHASE* (*CHS*), *CHALCONE ISOMERASE* (*CHI*), *CHALCONE ISOMERASE*-LIKE (*CHIL*), *FLAVANONE 3-HYDROXYLASE* (*F3H*), *FLAVONOID 3-HYDROXYLASE* (*F3’H*), and *FLAVONOL SYNTHASE 1* (*FLS1*) exhibit similar expression patterns in the root, with peaks of expression at the end of the MZ and in the DZ (fig.S6, A). Consistent with this, *MYB12,* which regulates the expression of these genes (*45*) (Fig.3), reaches its maximum expression level in the DZ (fig.S6, A). Also, as previously described in the analysis of the tissue-specific localization of these proteins (*52*), these genes appeared highly expressed in ground tissue and stele, but they were barely detected in the epidermis (fig.S6B) (*35*).

Because PSY1 phenotypes are observed in the EZ and DZ, we hypothesized that PSY1 may specifically regulate the biosynthesis of these secondary metabolites in these developmental zones. To test this, we analyzed *ProCHS:CHS-GFP* and *ProFLS1:FLS1-GFP* reporter lines (Fig.4, A and E). These lines express *CHS* and *FLS1* fused to GFP under the control of their native promoters. We found that synthetic PSY1 treatment increased GFP expression of both *ProCHS:CHS-GFP* and *ProFLS1:FLS1-GFP* in the DZ (Fig.4, B and F). No significant changes were observed at the end of the MZ or at the onset of root hair development (Fig.4, C, D, G, and H).

**Figure 4.**
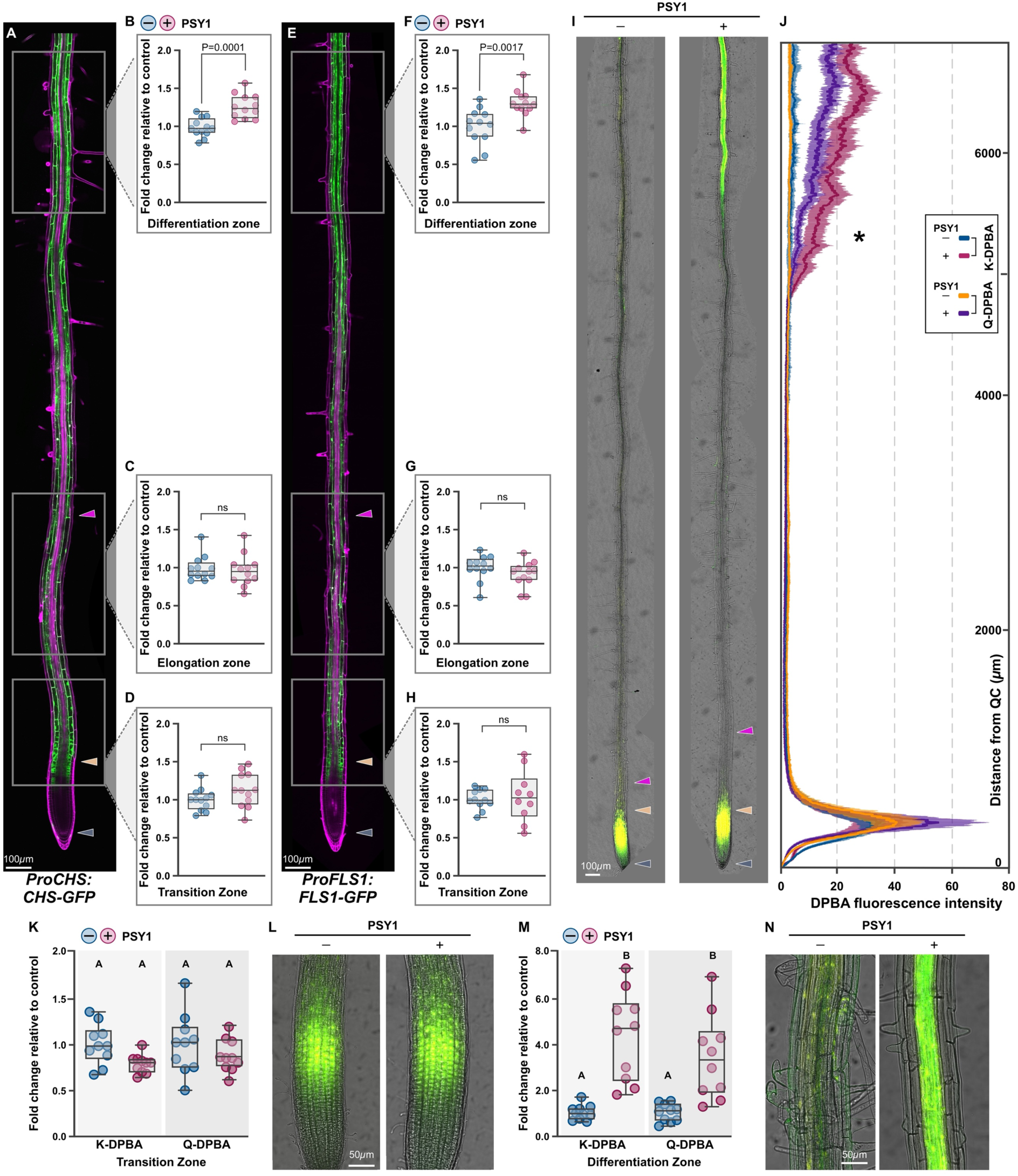
PSY1 induces the accumulation of flavonols in the differentiation zone. Expression of GFP reporter lines of *CHS* (*ProCHS:CHS-GFP*) (**A-D**) and *FLS1* (*ProFLS1:FLS1-GFP)* (**E-H**). The images in (**A**) and (**E**) show GFP fluorescence in green and cell walls stained with propidium iodide (PI) in magenta. In the regions marked by rectangles, *ProCHS:CHS-GFP* (**B**, **C**, and **D**) and *ProFLS1:FLS1-GFP* (**F**, **G**, and **H**) GFP intensity was quantified in 6-day-old seedlings transferred to 1xMS vertical plates with or without 100nM of synthetic PSY1 for 6 hours. Sum projections generated from 30 z-section images for each region were used for this quantification. (**I**) 5-day-old wt seedling roots were stained with DPBA (2-aminoethyl diphenylboric acid) after 24 hours with or without 250nM PSY1 treatment. DPBA is a probe that allows visualization of Kaempferol (green, K-DPBA) and Quercetin (yellow, Q-DPBA) flavonols. Images are scaled to correspond to the y-axis of the plot profile in (**J**). (**J**) Plot profiles of DPBA fluorescence in 5-day-old wt seedlings grown with or without PSY1 250nM for 24 hours. Data are mean ± s.e.m. of **n=12** seedlings. The asterisk at ∼5300μm from the QC represents the shortest distance at which the values for K and Q-DPBA fluorescence in the presence of PSY1 became significantly different (P<0.05) from the controls as determined by unpaired two-tailed Student’s t-test. The fluorescence intensity of K-DPBA and Q-DPBA was quantified in the transition zone (**K** and **L**) and in the differentiation zone (**M** and **N**) using sum projections generated from 30 and 50 z-section images, respectively, for each region. In (**B**), (**C**), (**D**), (**F**), (**G**), (**H**), (**K**), and (**M**), the fluorescence intensity is plotted as a fold change relative to the control. The data shown are box and whisker plots combined with scatter plots; each dot represents an independent seedling measurement. Representative images of two independent experiments (**n=10-12** seedlings/experiment) are shown. In (**B**), (**C**), (**D**), (**F**), (**G**), and (**H**), P-values are calculated by a two-tailed Student’s t-test. In (**K**) and (**M**), different letters indicate significant differences, as determined by one-way ANOVA followed by Tukey’s multiple comparison test (P < 0.05). The purple arrowheads mark the position of the QC, the pale orange arrowheads mark the end of the meristem, where cells start to elongate, and the magenta arrowhead points to the first root hair bulge, defined as stage +2, indicating the end of the elongation zone in the epidermis.

We next investigated whether CHS and FLS1 expression patterns correlate with metabolite accumulation. For these experiments, we utilized the flavonol-specific dye diphenylboric acid 2-aminoethyl ester (DPBA). Kaempferol-DPBA (K-DPBA) and quercetin-DPBA (Q-DPBA) exhibit distinct spectral properties, enabling independent quantification of these two flavonols (*53*). Consistent with the localization of flavonol biosynthetic enzymes, K-DPBA and Q-DPBA signals were detected both at the end of MZ and in the DZ (Fig.4I), with fluorescence peaking 300μm from the root tip (*52*). To visualize and quantify DPBA fluorescent signals along the root longitudinal axis, we generated plot profiles for K-DPBA and Q-DPBA fluorescence intensity from the QC to the DZ (Fig.4, I and J). In PSY1-treated roots, K-DPBA and Q-DPBA fluorescence became significantly stronger in the DZ, approximately 5300μm from the QC (Fig.4, I and J). Because the images used for these profiles were obtained using the middle focal plane of the root, we also measured K-DPBA and Q-DPBA fluorescence intensity using z-stacks covering the complete root width and observed a significant increase in DPBA signal in the DZ (Fig.4, M and N), while no change was evident in the TZ (Fig.4, K and L). These results indicate that PSY1 induces upregulation of genes encoding flavonoid biosynthetic enzymes and accumulation of flavonols in the DZ.

### 4. Flavonol biosynthesis is required for PSY1-induced root growth

We then explored whether PSY1 could enhance root growth in flavonoid-deficient mutants, also known as *transparent testa (tt)*, due to their seed coat color phenotype (*54*). We tested four out of 11 flavonoid pathway genes that were upregulated by PSY1 treatment in our RNA-seq dataset (Fig.3). The selection included Col-0 plants with mutations in early flavonoid biosynthesis steps: *tt4-11* (*chs*) and *tt5-2* (*chi*) that lack of flavonoids (*52*), as well as *tt7-7* (*f3′h*), which contains a T-DNA insertion in the gene encoding *F3′H* (*54*) and is known to accumulate only kaempferol. We also examined *myb12*, which carries a mutation in the *MYB12* transcription factor and does not accumulate flavonols in the root (*55*). We also utilized multiple single- and double-point mutant alleles available in the Ler genetic background, including *85* (*tt4*), *86* (*tt5*), *88* (*tt7*), and *8592* (*tt4tt7*) (*51*, *56*).

We then generated multiple independent transgenic lines expressing *Pro35S:PSY1* in wt and flavonol-deficient plants in the Col-0 genetic background (Fig.5 and fig.S7A). The Ler mutants were subjected to synthetic PSY1 treatment (fig.S8). We found that PSY1 ectopic expression or synthetic PSY1 treatment led to significantly longer primary roots only in wt seedlings (Fig. 5A; fig.S7E and fig.S8A). Under our growth conditions, the flavonoid biosynthetic enzyme mutants developed longer MZ compared to the wt, as previously observed by Silva-Navas and colleagues (*57*), with the exception of *85 (tt4)* and *8592 (tt4tt7)* in the Ler genetic background providing an example of how identical mutations can yield distinct phenotypes in different genetic backgrounds (fig.S7, B-D and fig.S8B). In line with our previous finding that PSY1 does not control the position of the TZ, ectopic PSY1 expression or synthetic PSY1 treatment did not affect the MZ length in both wt and flavonoid-deficient mutants (fig.S7, B-D and fig.S8B). To validate these results, we also compared MZ cell number in *Pro35S:PSY1 vs.* Empty Vector (EV, as control) in wt and *tt4-11* using a cortical cell length profile (fig.S7, B and C) and detected no significant changes in the MZ size. Moreover, we found that the cell elongation profiles were almost indistinguishable between *tt4-11* with ectopic PSY1 expression (*tt4-11-Pro35S:PSY1)* and *tt4-11* with EV. In contrast*, Pro35S:PSY1* expression in a wt background caused a significant increase in mature cell length (Fig.5,B and C, fig.S2D and fig.S7, B and F). In *tt4-11* and *8592 (tt4tt7)* mutants, mature cortical cells were shorter than in wt, and their size was less responsive to PSY1 expression (Fig.5, B and C; fig.S7F and fig.S8, C and D). Despite no significant differences in mature cortical cell length observed among other mutants compared to the wt, we successfully confirmed a consistently reduced response to both PSY1 overexpression and synthetic PSY1 treatment (Fig. 5, B and C; fig.S7F and fig.S8, C and D). These data suggest that PSY1-mediated control of mature cortical cell size relies on both kaempferol and quercetin accumulation.

**Figure 5.**
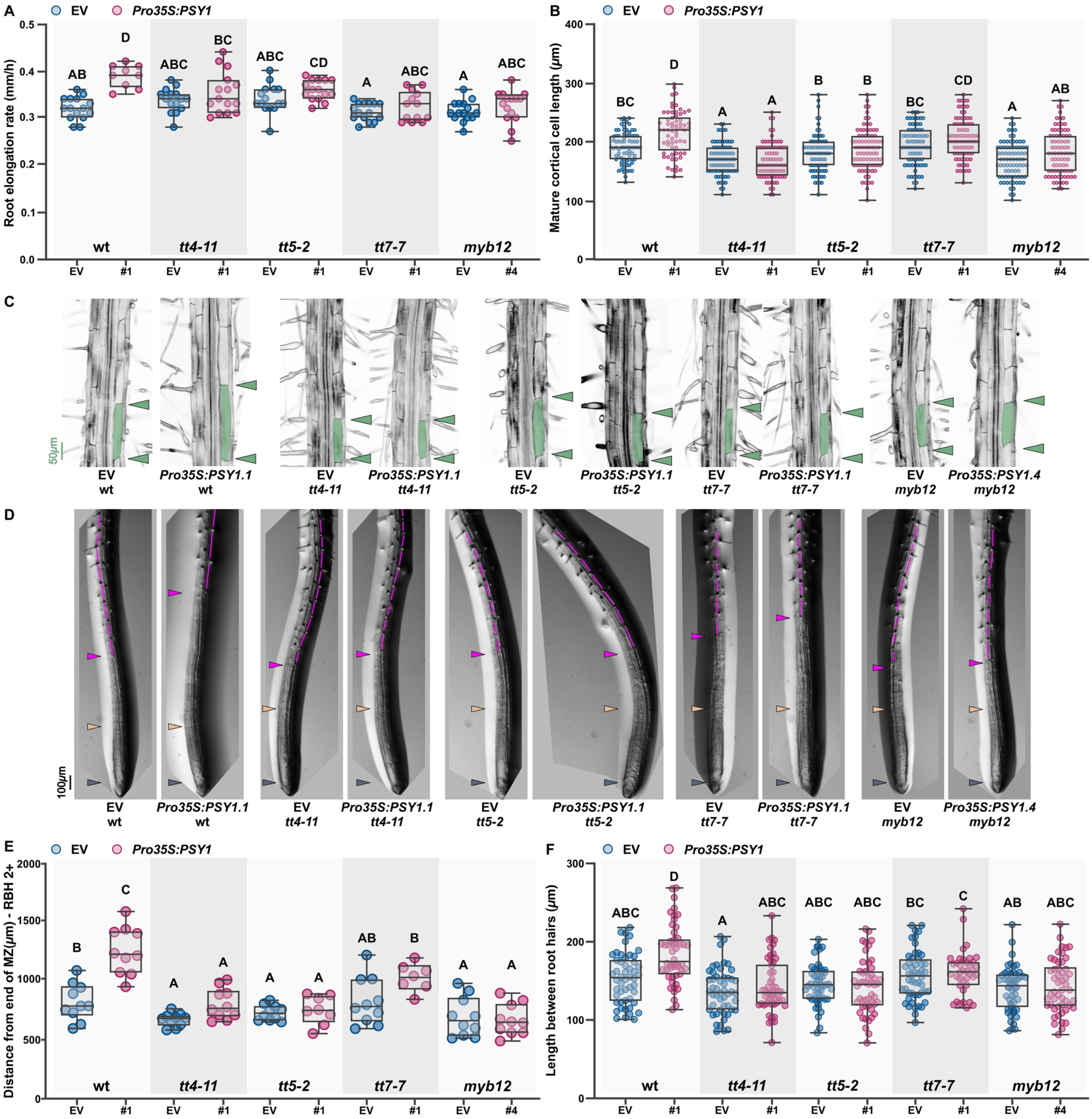
Flavonol biosynthesis is required for PSY1 control of root zonation. (**A**) Root elongation rate (mm/h)(**n=10-15** seedlings),(**B** and **C**) mature cortical cell length (**n=80** cells), (**D**) root tip architecture,(**E**) distance from the end of the MZ to first root hair bulge at stage +2 (RHB 2+) (**n=8-10** seedlings),(**F**) length between consecutive root hairs in one trichoblast file (**n=50**) in 7-day-old independent homozygous transgenic lines that accumulated higher levels of PSY1 (*Pro35S: PSY1*) with EV (empty vector) control generated in wild-type (wt-Col-0) and mutant plants defective in flavonol biosynthesis (*tt4-11, tt5-2,tt7-7 and myb12*). Only one homozygous transgenic line overexpressing *PSY1* is included in this figure; two more lines with similar results are presented in Supplemental figure 7. In (**C**), the limits of a representative mature cortical cell are shaded in green. In (**D**), the length between consecutive root hairs in one trichoblast file is highlighted in magenta. In (**A**), (**B**), (**E**), and (**F**), the data shown are box and whisker plots combined with scatter plots; each dot indicates the measurement of the designated parameter listed on the y-axis of the plot. Different letters indicate significant differences, as determined by one-way ANOVA followed by Tukey’s multiple comparison test (P < 0.05). The purple arrowheads mark the position of the QC, the pale orange arrowheads mark the end of the meristem, where cells start to elongate, the green arrowheads indicate the mature cortical cell size, and the magenta arrowhead points to the first root hair bulge, defined as stage +2, indicating the end of the elongation zone in the epidermis.

We also observed that in the *tt4-11* mutant, root hairs developed significantly closer to the end of the MZ compared with wt plants (Fig.5, D and E and fig.S7G). This phenotype was also observed in all flavonol-deficient mutants in the Ler background (fig.S8E). These findings align with previous studies indicating that *tt4-11* mutants exhibit a higher number of root hairs in a region closer to the TZ (*52*). Gayomba and Muday (*52*) also found that the length between consecutive root hairs in one trichoblast file was reduced in *tt4-11* compared to wt; we observed a similar trend. However, under our growth conditions, these results were not statistically significant (Fig.5, D and F and fig.S7H). Together, these results suggest that morphological signs of differentiation are shifted toward the root tip in flavonol-deficient plants, emphasizing the role of flavonol biosynthesis in proper root zonation. Additionally, when examining root hair initiation, we found that it occurred farther away from the root tip in wt plants expressing PSY1 ectopically or grown in media supplemented with synthetic PSY1. This significant response was not observed in other flavonoid mutants tested (Fig.5, D-F; fig.S7, G and H and fig.S8E).

Taken together, these results indicate that flavonol biosynthesis is required for PSY1-dependent regulation of root growth, supporting a model where flavonol accumulation acts downstream of PSY1 signaling.

### 5. Synthetic Naringenin treatment phenocopies PSY1 treatment

To test the role of flavonols in root zonation, we treated plants with Naringenin, a flavonoid precursor, and compared them with plants treated with PSY1. We hypothesized that increasing flavonol levels in the differentiation zone would mimic the effects of PSY1 overexpression.

First, we confirmed that Naringenin could be converted to different flavonol products in *tt4-11* roots by detecting the K-DPBA and Q-DPBA signals (fig.S9A). Next, we examined whether Naringenin could rescue root elongation defects in the *tpst-1* mutant as PSY1 does. For this experiment, we transferred 5-day-old *tpst-1* seedlings to plates with different concentrations of Naringenin (10, 25 or 50μM) or with PSY1 (50nM) and measured root elongation after 48 hours. We found that PSY1-treated plants had significantly longer primary roots than the untreated plants and that Naringenin also induced root growth at 25μM and 50μM (fig.S9B).

We then analyzed the cell length profile of *tpst-1* cortical cells from the QC to the DZ and observed a striking similarity between PSY1 and 25 μM Naringenin treatments (fig.S9, C and D). It has been previously shown that 8-day exposure to quercetin significantly reduced root meristem size in the wt background (*57*). However, we found that none of the treatments altered the size of the MZ in *tpst-1*, indicating that under these conditions, Naringenin did not affect the position of the TZ in this mutant background (fig.S9D). To examine how Naringenin affects cell elongation in *tpst-1* mutants, we measured the mature cortical cell length and the length between consecutive root hairs in one trichoblast file in *tpst-1* plants treated with 25 μM Naringenin. We found that both parameters increased, mimicking the effects of PSY1 treatment in this mutant (Fig.6, A-D and fig.S9, C and D). These results suggest that flavonols act downstream of the PSY1 signaling pathway, controlling the magnitude of cell elongation.

**Figure 6.**
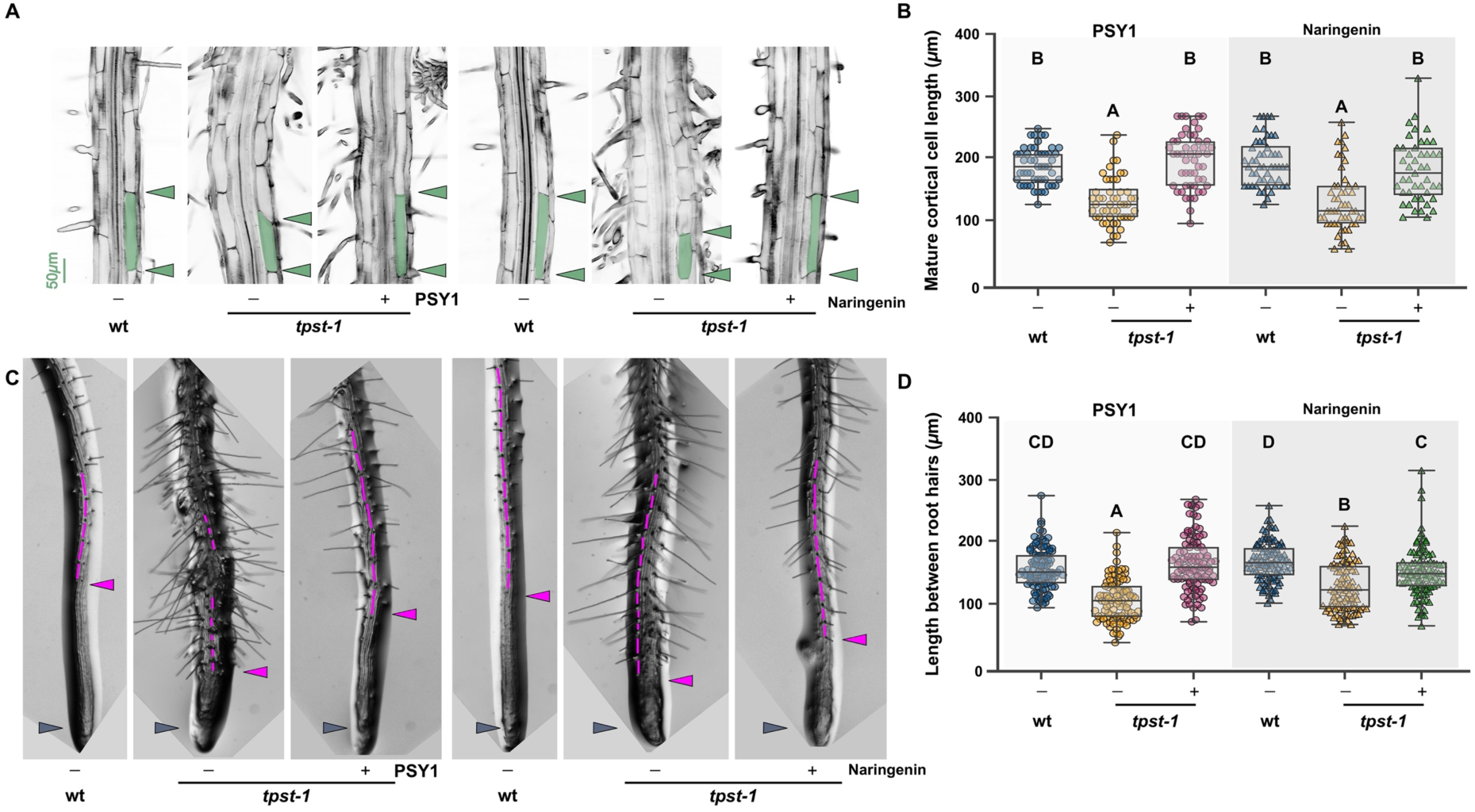
Exogenous treatment with Naringenin rescues cell expansion defects in *tpst-1.* (**A** and **B**) Mature cortical cell length (**n=50** cells), (**C**) root tip architecture, and (**D**) length between consecutive root hairs in one trichoblast file (**n=100** cells) in 5-day-old wt and *tpst-1* seedlings grown for 48 hours in control conditions (1xMS or 1xMS supplemented with EtOH, -) or treated with 50nM PSY (+) or 25μM Naringenin (+). In (**A**), the limits of a representative mature cortical cell are shaded in green. In (**C**), the length between consecutive root hairs in one trichoblast file is highlighted in magenta. In (**B**) and (**D**), the data shown are box and whisker plots combined with scatter plots; each dot indicates the measurement of the designated parameter listed on the y-axis of the plot. Different letters indicate significant differences, as determined by one-way ANOVA followed by Tukey’s multiple comparison test (P < 0.05). The purple arrowheads mark the position of the QC, the green arrowheads indicate the mature cortical cell size, and the magenta arrowhead points to the first root hair bulge, defined as stage +2, indicating the end of the elongation zone in the epidermis.

### 6. Changes in PSY1 signaling alter auxin activity and H_2_O_2_ accumulation

Our results indicated that PSY1-induced root growth requires flavonol biosynthesis in the DZ. We next looked for potential flavonol targets involved in this process. Flavonols are known to affect root growth through two mechanisms: regulation of polar auxin transport (PAT) through inhibition of PIN-mediated auxin efflux and maintenance of reactive oxygen species (ROS) homeostasis (*58*). Based on these reports, we assessed whether increased levels of PSY1 altered either of these processes.

We examined the effect of PSY1 on auxin signaling in the root stele with the *DR5v2:3nGFP* reporter (Liao et al., 2015). Consistent with our previous results, synthetic PSY1 did not alter auxin activity in the meristem after 6-day treatment (Fig.7, A and B and fig.S4, E and F). However, PSY1 reduced *DR5v2:3nGFP* activity in the stele, starting 1500μm away from the QC (Fig7, A and B). Because high auxin levels can cause cell wall alkalization and inhibit cell elongation (*59*, *60*), we hypothesize that the reduction of auxin activity in the EZ/DZ caused by PSY1 could signal cells to continue to elongate resulting in significantly longer mature cell sizes compared to those of wt plants.

**Figure 7.**
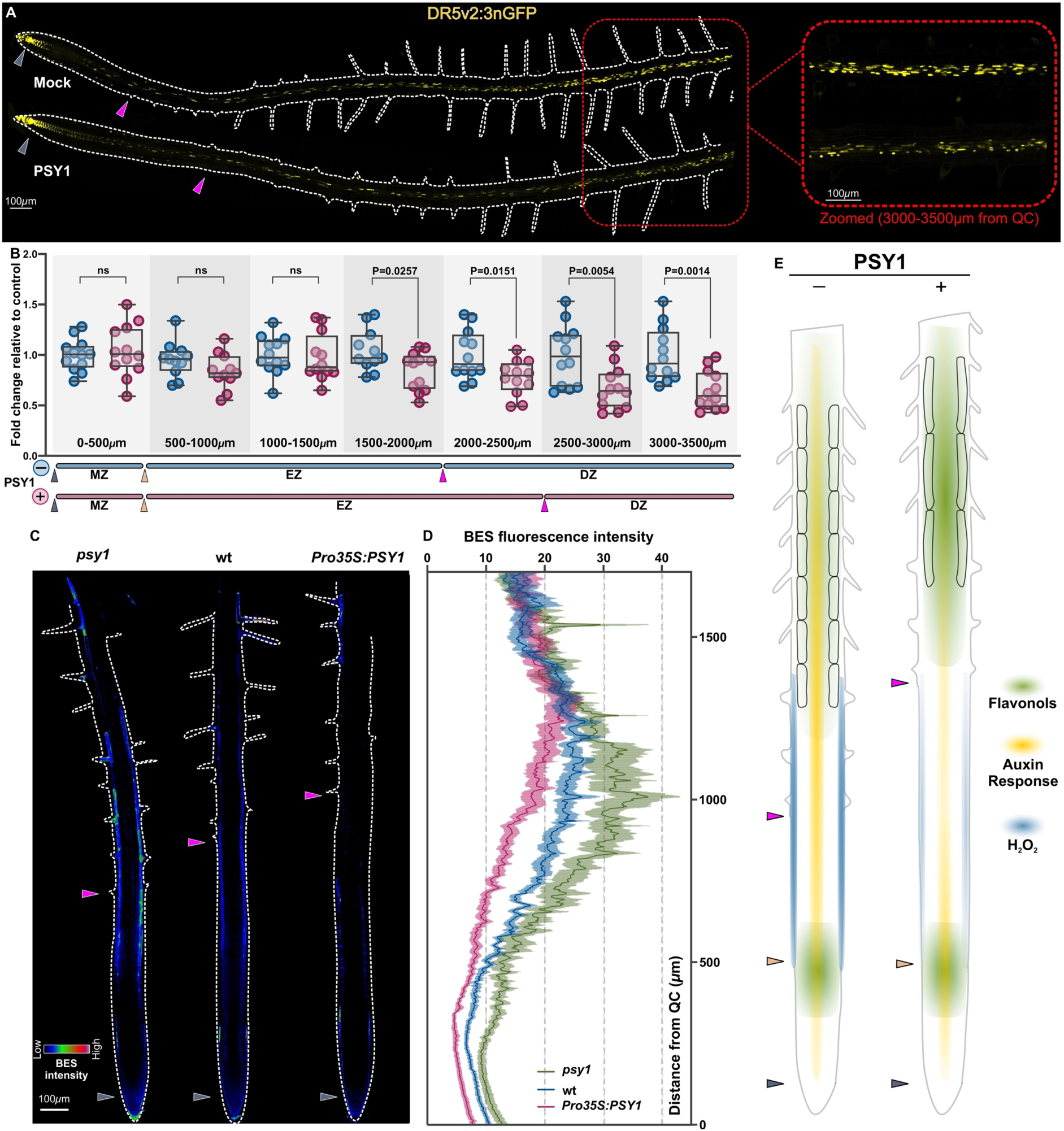
PSY1 treatment alters auxin activity in the stele and H_2_O_2_ accumulation in the epidermis. (**A**) Activity of the auxin response reporter line, *DR5v2:n3GFP* (yellow) in 6-day-old roots grown in the presence or absence of 100nM PSY1. (Left) Longitudinal cross-sections of the roots; organ boundaries are marked by white dashed lines. (Right) Magnified images of *DR5v2::n3GFP* expression in the stele in the differentiation zone are highlighted by a red box (3000-3500μm from the QC). (**B**) Changes in *DR5v2::n3GFP* expression in response to PSY1 treatment in the stele along the root longitudinal axis from the QC (0μm) to the differentiation zone (3500μm). The sizes of the different developmental zones measured in the root cortex are depicted in blue and pink for plants grown without and with PSY1, respectively. In (**B**), fluorescence intensity is plotted as a fold change relative to the control. The data shown are box and whisker plots combined with scatter plots; each dot represents an independent seedling measurement. Representative images of four independent experiments (**n=10** seedlings/experiment) are shown. P-values are calculated by a two-tailed Student’s t-test. (**C**) Representative epidermal H_2_O_2_ accumulation along the root longitudinal axis using BES-H2O2-Ac (BES) of *psy1*, wt, and *Pro35S:PSY1* 6-day-old seedlings. BES fluorescence intensity profile is shown using RGB-rainbow false color with its color scale bar; organ boundaries are marked by white dashed lines. In (**C**), images are scaled to correspond to the y-axis of the plot profile in (**D**). (**D**) Plot profiles of epidermal BES fluorescence in *psy1*, wt, and *Pro35S:PSY1* 6-day-old seedlings. Data are mean ±s.e.m. of **n=15** seedlings. (**E**) A diagram summarizing the observed effects of increased levels of PSY1 in the root. Accumulation of PSY1 is associated with activation of flavonol biosynthesis in the DZ, reduction of auxin responses in the stele of the DZ, and reduction in H_2_O_2_ in the epidermis of EZ/DZ. Furthermore, morphological changes associated with cell differentiation, such as root hair initiation, are observed further away from the root tip of plants that accumulate high PSY1. The purple arrowheads mark the position of the QC, and the magenta arrowheads mark the end of the elongation zone in epidermis defined by the appearance of the root hairs.

We next assessed possible changes in ROS quantity or distribution in response to PSY1 signaling using nitro blue tetrazolium (NBT) staining to detect O^2-^ and BES-H_2_O_2_-Ac fluorescence to detect H_2_O_2_ (*40*). We used RGF1-treated plants as a control because RGF1 is known to alter ROS accumulation to control MZ size (*40*). RGF1 increased total NBT intensity in the MZ (fig.S10, C and D), but loss or ectopic expression of PSY1 in *psy1* or *Pro35S:PSY1,* respectively, did not affect NBT intensity (fig.S10, A-D). These results provide further evidence that PSY1 does not regulate MZ size in a ROS-dependent manner. It has been proposed that flavonol modulation of ROS accumulation is one of the mechanisms driving root hair initiation (*52*). We, therefore, hypothesized that PSY1 signaling may affect root hair development by modulating H_2_O_2_ levels in the root epidermis. To test this, we measured H_2_O_2_ accumulation in the epidermis along the root longitudinal axis using BES-H_2_O_2_-Ac. We compared BES-H_2_O_2_-Ac fluorescence intensity in wt, *psy1*, *Pro35:PSY1,* and RGF-treated plants. RGF1 treatment led to a longer MZ with lower BES-H_2_O_2_-Ac fluorescence intensity compared to the untreated control (fig.S10, E and F)(*40*). In contrast, we found that *Pro35:PSY1* plants had lower H_2_O_2_ levels than wt plants, while *psy1* plants had higher H_2_O_2_ levels, although the overall H_2_O_2_ epidermal profile remained the same as wt (Fig.7, C and D and fig.S10G). These results suggest that PSY1 signaling negatively regulates H_2_O_2_ production in the root epidermis, possibly influencing root hair initiation.

A diagram summarizing the observed effects of increased levels of PSY1 in the root is shown in Fig.7E.

## Discussion

Root zonation is the spatial arrangement of the cells along the root longitudinal axis reflecting the balance between cellular growth and maturation (Fig.1A, left side of the panel). The sustained growth of the root requires these processes to be tightly coordinated. Previous studies have shown that several phytohormone and peptide signaling pathways regulate the establishment of boundaries that separate cells in different developmental stages in the root, therefore controlling root zonation (*12*, *61*). The tyrosine sulfated peptide hormone family known as PSY has been shown to participate in the control of cortical mature cell size in Arabidopsis primary roots (*29*, *30*). Despite recent progress in characterizing the LRR-RLKs involved in peptide perception, the biological processes triggered by PSYs remain largely unexplored. In this study, we focused on one of the members of this family, PSY1. We started by investigating the function of PSY1 in root zonation based on a detailed cell length profile analysis (Fig.1D). We observed that *psy1* mutants have shorter mature cells in both the root cortex and epidermis, leading to reduced root growth (Fig.1H and K). Additionally, we found that defining mature cell features, such as root hair initiation and deposition of secondary cell wall in the protoxylem, which marks the cessation of rapid cell elongation (*41*, *42*), emerged closer to the root tip in *psy1* plants than in wt plants (Fig.1, I-M). These phenotypes could be reversed when synthetic PSY1 is supplemented exogenously (Fig.1, I-M). Overall, PSY1 signaling regulates root growth by modulating the magnitude that cells elongate before reaching their final, differentiated size (Fig.7E). Given that our analysis is based on cell length, without specific data on growth rates, the exact effect of PSY1 on these observed phenotypes remains to be fully understood. It could be due to PSY1 signaling (1) controlling the maximum cellular growth rate, which is the speed at which cells elongate before leaving the rapid growth region; (2) determining the onset of growth cessation, or the position at which the cellular growth rate began to decrease shootward in the EZ; or (3) influencing a combination of both. Finally, it is worth noting that the overall progression to the root maturation was affected by PSY1 signaling, including tissues such as the epidermis where the expression of PSY1 promoter was barely detected (Fig.1A and fig.S1, B and D). This observation aligns with the nature of these peptides being secreted and diffusible, potentially creating a gradient from their site of synthesis. Consequently, the expression pattern of ProPSY1-GFP might not accurately represent the regions where PSY1 is active and perceived by its extracellular receptors.

The role of PSY1 in controlling developmental transitions was previously described at the onset of the embryo-to-seedling transition, which is associated with the establishment of a well-sealed cuticle required for an aerial lifestyle. In the model proposed by De Giorgi and colleagues (*33*), tyrosine-sulfated peptides, including PSY1, are released towards the embryo as part of the endosperm secretome to signal the formation of the seedling cuticle. In agreement with this model, *psy1* and endosperm-less seedlings had higher toluidine blue O uptake in cotyledons, suggesting cuticle defects in these plants (*33*). Transcriptomic analysis revealed that in the presence of light, GO categories such as flavonoid and glucosinolate biosynthetic processes were enriched among repressed genes in endosperm-less seedlings relative to wt (*33*). If we consider the lack of endosperm as a proxy for reduced PSY1 signaling in the developing seedling, this analysis supports a role for PSY1 in controlling the onset of the embryo-to-seedling transition through the expression of genes involved in secondary metabolite biosynthesis.

The transcriptomic data generated in this study revealed that Arabidopsis seedling roots treated with synthetic PSY1 exhibited increased expression of genes in the phenylpropanoid and flavonoid biosynthetic pathways (Fig.3 and Supplemental Data Set 1), suggesting that PSY1 activates these secondary metabolic pathways. The triple PSY receptor mutants, *tri-1*, develop longer roots, as observed with PSY1 overexpression, and also exhibit increased expression of genes that encode flavonol biosynthetic enzymes (fig.S5E)(*30*, *31*). It is worth noting that more flavonol pathway genes are differentially expressed in the triple receptor mutant compared to our synthetic PSY1 treatment. For example, in the *FLAVONOL SYNTHASE* (*FLS*), *4-COUMARATE: COA LIGASE* (*4CL*), and *PHENYLALANINE AMMONIA-LYASE* (*PAL*) gene families, only one isoform was activated by PSY1 treatment, while multiple isoforms were upregulated in the triple receptor mutant. Also, *O-METHYLTRANSFERASE 1* (*OMT1*), involved in the methylation of flavonols to generate isorhamnetin (*43*), for which we did not find changes after PSY1 synthetic treatment, is activated in the triple receptor mutant background, indicating that PSY genes may also regulate the accumulation of isorhamnetin scaffolds. The KEGG pathway analysis of the triple PSYR mutant also revealed enrichment for genes that participate in glucosinolate biosynthesis, including glucosinolates derived from methionine and aromatic amino acids, which were not detected in the PSY1 treatment (fig.S5E). The crosstalk between glucosinolate biosynthesis and flavonols has been recently explored by Naik and colleagues (*62*) as part of an effort to understand the molecular mechanisms underlying the ability of flavonols to control plant development. Transcriptomic and targeted metabolomic analysis hint at flavonols promoting the accumulation of aliphatic glucosinolates (*62*). Although our study did not assess changes in glucosinolate biosynthetic enzymes, it would be interesting to test if PSY1 affects this metabolic pathway using root samples generated after longer PSY1 synthetic treatment or in the *psy1* mutant.

It was recently shown that CLE-LIKE6 (CLEL6), a member of a different family of small tyrosine-sulfated peptides, inhibits the biosynthesis of anthocyanin, another type of flavonoid (*63*). *CLEL6* expression in the hypocotyl decreases during photomorphogenesis, which activates anthocyanin biosynthesis genes and leads to pigment accumulation. These pigments help regulate ROS levels and facilitate seedling development during de-etiolation (*63*). The contrasting effects of PSY1 and CLEL6 may be due to tight spatial control of flavonoid enzyme gene expression, which allows only flavonols, not anthocyanins, to accumulate in roots. In support of this idea, we found that PSY1 upregulates MYB12, a flavonol-specific flavonoid regulator (*45*), but does not affect *PRODUCTION OF ANTHOCYANIN PIGMENT1 (PAP1)*, which controls late anthocyanin biosynthesis genes (Fig.4 and Supplemental Data Set 1) (*64*).

The results presented in this paper shed light on how flavonols accumulate in a developmental zone-specific manner during root development. One of the few examples of a root growth modulator affecting the flavonoid pathway is the transcription factor WRKY23 (*65*). Specifically, WRKY23 induces *FLAVONOID 3-HYDROXYLASE* (*F3’H*) expression to negatively influence auxin transport from the shoot to the root. Overexpression of WRKY23 leads to higher flavonol levels in the whole root, including TZ and DZ, reducing rootward auxin transport and causing root tip disorganization. Accumulation of flavonols in the TZ has also been studied in detail by Silva-Navas et al., 2016. In this case, flavonols reduce auxin activity in the MZ and decrease *PLT* gene expression, providing a mechanism to explain their role as inhibitors of root growth. Given that PSY1-induced root growth requires the activation of flavonoid biosynthetic enzymes and flavonol accumulation, specifically in the DZ as detected by DPBA staining, we surmised that the effect of flavonols on root growth depends on where they accumulate within the root. To test this hypothesis, we used *tpst-1*, a mutant in which synthetic flavonol treatment does not modify MZ size. Because *tpst-1* plants have a shorter MZ (*40*, *66*), as a result of decreased PLT protein stability, it is possible that any additional increase in flavonol levels will not further inhibit meristematic activity. Complementation of cell elongation defects in *tpst-1* by Naringenin likely reflects the effects of flavonols in the DZ, phenocopying the effects of the PSY1 signaling pathway (Fig.6 and fig.S9, C and D).

Auxin activity measurement using a response reporter line (*DR5v2:3nGFP)* showed a marked difference at 1500μm from the quiescent center (QC) between PSY1-treated and untreated roots. This point coincides with the location where cortical cells of untreated plants stop growing, whereas cells continue to elongate in PSY1-treated plants (Fig.1D; fig.S2D and fig.S3D). It is possible that flavonol accumulation in the DZ could block auxin transport from the shoot, resulting in reduced auxin activity in the EZ. Moreover, the extended elongation phase observed in plants with high PSY1 levels might be due to a delayed onset of growth cessation. This delay could be linked to the role of cytokinin altering cell wall properties, which is dependent on an increase of auxin levels in the EZ (*9*, *67*).

The results presented here also suggested other rich areas for research. For example, two interesting pathways that appeared enriched among the downregulated KEGG included “MAPK signaling pathway” and “plant hormone signal transduction” (Fig.3E and Supplemental Data Set 1). The transcripts related to these pathways included *PYL4*, *PYL5*, and *PYL6*, which are members of the *PYRABACTIN RESISTANCE1* (*PYR1*) / *PYR1-like* (*PYL*) / *REGULATORY COMPONENTS OF ABA RECEPTOR* (*RCAR*) family of proteins involved in ABA perception (*68*). Also, the *1–AMINOCYCLOPROPANE-1–CARBOXYLATE (ACC) OXIDASE 1* (*ACO1*), which oxidases ACC to ethylene (*69*), was downregulated. It is known that ABA and ethylene signals are integrated to mediate root growth inhibition (*70*). PSY1 suppression of these hormonal pathways is consistent with the shootward displacement of differentiation signs. Thus, genetic interactions between PSY1 and ABA and ethylene pathways can be assessed for their effects on the transition from active growth to differentiation.

The PSY-PSYR signaling pathway has been implicated in mediating the trade-off between growth and stress responses in various plant species (*30*). However, the specific roles of each PSY peptide and PSYR remain largely unknown. Our results show that PSY1 acts as a repressor of growth cessation through modulation of flavonols, which are known to control plant growth and stress responses. Given that flavonols act as ROS scavengers (*71*, *72*), increased accumulation of flavonols downstream of the PSY1 signaling may, therefore, explain the reduction of H_2_O_2_ levels in the EZ. Furthermore, we observed a shootward displacement in two cell maturation processes that rely on ROS levels in plants that ectopically accumulate PSY1: root hair development and lignin deposition (*18*, *52*) (fig.S2, I-M and Fig.6, C and D). Additionally, it is well-established that ROS can help defend the cell against invading bacterial and viral pathogens (*73*). Thus, higher levels of PSY1, leading to increased accumulation of flavonols, would serve to reduce H_2_O_2_ in the tissue where PSY1 is expressed, suggesting a role for PSY1 in mediating the trade-off between growth and stress responses (Brunetti et al., 2018; Lee et al., 2020). Thus, we hypothesize that PSY-rich tissues would be more susceptible to pathogen infection. Support for this hypothesis is reflected in the observation that the rice bacterial pathogen *Xanthomonas oryzae pv. oryzae* (*Xoo*), which produces a molecular mimic of PSY1 named RaxX (Required for activation of XA21-mediated immunity X), showed reduced virulence in the absence of RaxX (*32*, *74*). Like PSY1, RaxX can promote root growth and can bind to the PSYR in Arabidopsis (*30*, *32*). These observations suggest that RaxX, mimicking a growth-promoting peptide hormone, may modify the developmental processes in a way that favors bacterial infection (*24*). Future experiments directed at assessing the accumulation of antioxidant flavonols in leaf vascular tissues and the effect on plant susceptibility to *Xoo* infection would help address this question.

## Materials and Methods

### Plant Materials, Growth Conditions, and Treatments

*Arabidopsis thaliana* accession Col-0 was used throughout this study as the wild-type (wt) background, unless otherwise indicated. See Supplemental Tables 1 and 2 for a full list and description of the mutants and reporter lines utilized in this study. Seeds were surface sterilized using 70% ethanol for 10 min and then rinsed three times with absolute ethanol. Seeds were stratified in 0.1% agarose at 4°C for three days before germination. Plates were prepared with standard MS medium (1X Murashige and Skoog salt mixture with vitamins, MSP09-Caisson Laboratories), 1% sucrose, and 0.3% gellan gum (G024-Caisson-Gelzan) and adjusted to pH 5.65 with KOH. Seeds were placed on the plates (20 seeds per plate), and the lids were secured with Micropore surgical tape (1530-0). Seedlings were grown in vertically positioned plates in a chamber with long photoperiods (16 h light/8 h dark) at 21°C. Germination rate was scored in every experiment, and no significant differences between genotypes and treatments were observed.

For synthetic peptide treatments, peptide (or water for untreated plants) was added to the MS media before pouring it into a plate. The length and concentration of the peptide treatment are specified in the figure legends. Seedlings were either germinated on media or moved after germination to treatment plates. All peptides used in the experiments are tyrosine sulfated. The synthetic PSY1 peptide lacks the hydroxy- and L-Ara3-modifications at the C-terminus and was obtained from Pacific Immunology (Ramona, CA, USA). RGF1 was obtained from Peptide 2.0 (Chantilly, VA, USA). Peptides were diluted in ddH_2_O to a final concentration of 1mM. For synthetic flavonoid treatments, plants were grown on 1X MS media prepared as described above for 5 days and subsequently moved to 1X MS medium containing different concentrations of Naringenin or ethanol (mock treatment) for 48 hours. Stock solutions of Naringenin (Indofine Chemical Company) were freshly prepared to a final concentration of 100mM in absolute ethanol. For each experiment, MS media was freshly prepared and cooled for 1 hour in a 55-60°C water bath after autoclaving before adding chemicals.

### Cloning and Generation of Transgenic Lines

DNA constructs were created with the Gateway cloning technology (*75*). The genomic PSY1 sequence and the 1200pb PSY1-promoter, including the 5’UTR, were amplified using the primers described in Supplemental Table 3. These sequences were then recombined with pENTR^TM^/D-TOPO (Invitrogen, Cat#45-0218) to yield pTOPO_PSY1 and pTOPO_ProPSY1. The latter vectors were used in a Gateway LR cloning (Gateway® LR Clonase^TM^ II Plus Enzyme Mix, Invitrogen; Cat#:12538-120) with pEarleyGate100 (*76*) and pGWB504 (*77*) to yield *Pro35S:PSY1* and *ProPSY1:GFP* constructs. The generated vectors were transferred to Agrobacterium tumefaciens strain GV3101, which was used in floral dip transformations. *Pro35S:PSY1* and pEarleyGate100 (Empty Vector-EV) transformants were obtained in Col-0, *tt4-11*, *tt5-2*, *tt7-7*, *myb12,* and Col-0 expressing the construct *pCYCB1;1:GFP*. *ProPSY1:GFP* lines were generated in Col-0.

### Analysis of Root Growth

For root elongation measurements, seedlings were grown vertically for six to seven days, depending on the experiment. Starting from day three after sowing until the end of the experiment, a dot was drawn at the position of the root tip. Finally, plates were photographed, and the root length was measured over time with Fiji Is Just ImageJ (*78*). Root growth rate, expressed in millimeters per hour, was estimated from root length (millimeters) vs. plant age (days after sowing) plots.

Six or seven-day-old seedlings were imaged under a bright field using a Zeiss Discovery 20 equipped with an Axiocam 506 color camera. The picture was taken to highlight the appearance of the first root hair bulge, defined as the first observed root hair in stage +2 (*20*). In addition, the distance between root hairs was assessed in a continuous file of epidermal trichoblasts cells using Fiji.

### Confocal Microscopy

Laser scanning confocal microscopy (LSCM) was performed throughout the study using a Plan Apochromat 20x/0.75 CS2 lens on a Leica TCS SP8 microscope. For all the reporter line analyses, roots were stained with 15 μg/mL propidium iodide (PI) (Sigma), rinsed, and mounted in water, except for *pTCSn::GFP*, in which plants were directly mounted without staining. Fluorescence signals were visualized after excitation by a 488-nm laser line for GFP, YFP, and PI or by a 448-nm laser line for CFP. The fluorescence emission was collected between 600 and 700 nm for PI, 495 and 555 nm for GFP and YFP, and 465 and 570 nm for GFP. Fluorescence intensity measurements for *ProPLT1:CFP* and *ProPLT1:PLT1:YFP* were performed as described in (*79*). For *ProPSY1:GFP*, *ProCHS:CHS-GFP*, *ProFLS1:FLS1-GFP*, *pTCSn::GFP,* and *DR5v2::n3GFP*, sum intensity projections were generated from 30 z-section images taken in different locations along the root longitudinal axis. After removing the background, the average GFP intensity was measured, and values were expressed as a fold change relative to the control plants. The fluorescence signal was only measured in the stele for the auxin response reporter line *DR5v2::n3GFP*. *pTCSn::GFP* seedlings were five days post-germination at the time of the transfer to plates containing either mock or PSY1 synthetic peptide. The gain for GFP acquisition in *ProPSY1:GFP* analysis was set to avoid saturation in the differentiation zone.

For combined cell wall and lignin staining, seedling fixation and staining were performed using an adapted Clearsee protocol (*80*). Briefly, six or seven-day-old seedlings were fixed for 1h at room temperature in 10% neutral buffered formalin in PBS, using 6-well plates, then washed five times for 1min with PBS 1X. Once fixed, seedlings were cleared in Clearsee solution for at least 24 hours under mild shaking. Fixed and cleared samples were incubated overnight in a Clearsee solution supplemented with 0.2% Basic Fuchsin and 0.1% Calcofluor White. After 12 hours, the staining solution was removed, and samples were rinsed once in fresh Clearsee solution, then washed twice for at least 120 minutes in a renewed Clearsee solution with gentle shaking. Roots were carefully placed on a microscope slide with ClearSee and covered with a coverslip. Excitation and detection windows were set as follows: Basic Fuchsin excitation at 552 nm and detection between 600 and 650 nm, Calcofluor white for excitation at 405 nm, and detection between 415 and 570 nm. Central longitudinal section images were acquired to generate a cell length profile by measuring the length of every consecutive cortical cell located from the QC until the differentiation zone for each plant. Meristematic zone length is defined as the region of isodiametric cells from the QC up to the cell that was twice the length of the immediately preceding cell and was determined according to the file of cortical cells (*81*). Mature cortical cell length was assessed in 10 consecutive cells, starting six cells above the cortical cell closest to the epidermal cell with the first root hair bulge (*10*).

### DPBA staining

We analyzed flavonols accumulation in roots using the probe DPBA as described in (*53*), which allows for distinct visualization of kaempferol DPBA (K-DPBA) and quercetin DPBA (Q-DPBA) using LSCM. Briefly, individual seedlings were stained in 0.25% w/v DPBA (Sigma-Aldrich), which was dissolved in 0.01% Triton-X (v/v) in water on a rotary shaker at low speed for 7 minutes. The roots were washed in deionized water for 7 minutes on the same shaker and mounted in deionized water for imaging. Fluorescence signals were visualized after excitation by a 448 nm laser line, and the emission spectra were captured between 475-504 nm for K-DPBA and 577-619 nm for Q-DPBA. The specificity of the signal was tested using *tt4-11*, which doesn’t produce flavonols, grown with and without synthetic Naringenin, the precursor that *tt4-11* is unable to synthesize (fig.S9). All the images were acquired using identical settings, except for the laser intensity and digital gain that were increased for imaging in the differentiation zone. To generate the plot profiles in Fig.4 J, in which all developmental zones are imaged together, the gain was set to avoid saturation in the TZ and differentiation zone. Sum intensity projections were generated from about 30 z-section images taken in different locations along the root longitudinal axis. After removing the background, the average intensity of K-DPBA and Q-DPBA was measured, and values were expressed as a fold change relative to the control plants.

### Reactive oxygen species detection

Reactive oxygen species quantification was performed as described in (*40*). Briefly, for superoxide anion (O_2_^-^) quantification, seven-day-old seedlings were stained for 15 minutes in a solution of 200μM NBT in 20 mM phosphate buffer (pH 6.1) in the dark and rinsed twice with distilled water. Images for NBT staining were obtained using a 20X objective using a Leica DFC7000 T Camera. The total intensities of NBT staining in the meristematic zone were measured using Fiji (*78*). To detect hydrogen peroxide, we incubated 6-day-old seedlings with H_2_O_2_-3′-O-acetyl-6′O-pentafluorobenzenesulfonyl-2′-7′-difluorofluorescein-Ac (BES-H_2_O_2_-Ac) (*82*) (WAKO) 50μM for 30 min in the dark, then mounted them in 10 mg ml^−1^ PI in water. Roots were observed using a 20x objective with the LSCM. Excitation and detection windows were set as follows: BES-H_2_O_2_-Ac, excitation at 488 nm and detection at 500–550 nm; PI staining, excitation at 488 nm, and detection at 600-700 nm. Central longitudinal section images were acquired to quantify BES-H_2_O_2_-Ac in the root epidermis from the QC until the differentiation zone. The BES-H_2_O_2_-Ac intensity as a plot profile for fifteen seedlings was generated as described in (*79*) and averaged for final representation.

### Total RNA extraction and library preparation

Col-0 seedlings were grown on a permeable membrane placed on a clear agar MS. Five-days post-germination, seedlings were transferred to a control medium or medium containing PSY1 (50nM) by moving the membrane to the selected media. After 4 hours, the root tip from each seedling was dissected using an ophthalmic scalpel. For each treatment, three replicates of 200 root sections were generated. We extracted total RNA from the samples using Spectrum Plant Total RNA kit (Sigma). RNA samples were treated with DNase I during RNA extraction. RNA quality was examined using a 2100 Bioanalyzer (Agilent). The concentration of total RNA was measured by a Qubit (Invitrogen) instrument. mRNA was isolated from an input of 1000 ng of total RNA with oligo dT magnetic beads and fragmented to 300 bp - 400 bp with divalent cations at a high temperature. Using TruSeq stranded mRNA kit (Illumina), the fragmented mRNA was reverse transcribed to create the first strand of cDNA with random hexamers and SuperScript™ II Reverse Transcriptase (Thermo Fisher Scientific) followed by second strand synthesis. The double-stranded cDNA fragments were treated with A-tailing ligation with JGI’s unique dual indexed adapters (IDT) and enriched using 8 cycles of PCR. The prepared libraries were quantified using KAPA Biosystems’ next-generation sequencing library qPCR kit and run on a Roche LightCycler 480 real-time PCR instrument. Sequencing of the flowcell was performed on the Illumina NextSeq500 sequencer using NextSeq500 NextSeq HO kits, v2, following a 2x151 indexed run recipe.

### Differential expression analysis after PSY1 treatment

Raw fastq file reads were filtered and trimmed using the JGI QC pipeline, resulting in the filtered fastq file (*.filter-RNA.gz files). Using BBDuk (https://sourceforge.net/projects/bbmap/), raw reads were evaluated for artifact sequence by kmer matching (kmer=25), allowing 1 mismatch, and detected artifact was trimmed from the 3’ end of the reads. RNA spike-in reads, PhiX reads, and reads containing any Ns were removed. Quality trimming was performed using the phred trimming method set at Q6. Finally, following trimming, reads under the length threshold were removed (minimum length 25 bases or 1/3 of the original read length - whichever is longer). Filtered reads from each library were aligned to the TAIR10 Arabidopsis genome using HISAT2 version 2.1.0 (*83*). featureCounts (*84*) was used to generate the raw gene counts using gff3 annotations. Only primary hits assigned to the reverse strand were included in the raw gene counts (-s 2 -p --primary options). EdgeR (version 3.30.3)(*85*) was subsequently used to determine which genes were differentially expressed between pairs of conditions. Genes with a false discovery rate (FDR)-adjusted P value less than or equal to 0.05 were regarded as differentially expressed between the PSY and mock treatment. The gene expression data for the longitudinal and radial roots was obtained from (*34*).

An in-house R script was developed to generate heatmaps based on this data. The script utilized the Elbow and Silhouette methods to determine the appropriate number of clusters. After analyzing the data using both methods, we identified that the optimal number of clusters is four.

The enriched Gene Ontology (GO) groups among differentially expressed genes were identified using Panther DB (*86*), while the Database for Annotation, Visualization, and Integrated Discovery (DAVID)(*87*) (https://david.ncifcrf.gov/) was used for the Kyoto Encyclopedia of Genes and Genomes (KEGG) pathway enrichment analysis. For the comprehensive analysis of single-cell gene expression across specific time zones and cell types (*35*), expression data was sourced from the Arabidopsis Root Virtual Expression eXplorer (ARVEX), accessible at https://shiny.mdc-berlin.de/ARVEX/. For each gene of interest, data was extracted to pinpoint both the temporal zone and cell type specificity. To determine the average normalized gene expression value, we averaged the expression levels of each gene within the designated time zones and cell types. Furthermore, we quantified the extent of gene expression in each cell population. This was achieved by calculating the proportion of cells expressing a particular gene, which divided the number of cells exhibiting expression by the total cell count. GO, KEGG, and single-cell RNAseq data visualization was performed using a bubble plot generated utilizing the ggplot2 package in RStudio (version 2023.09.1+494) with R version 3.3.0+ (*88*). In the bubble plots for GO and KEGG, the y-axis shows the False Discovery Rate (FDR) in a negative Log10 scale, whereas the x-axis is fixed, and terms from the same KEGG/GO subtree are located closer to each other. The size of each circle represents the term Fold Enrichment in the Log10 scale.

### Gene Expression Analysis via RT-qPCR

Total RNA (1μg) was extracted from whole seven-day-old seedling tissue using TRIzol reagent (Invitrogen) and treated with the TURBO DNA-free kit (Ambion) to remove residual genomic DNA. cDNA was synthesized using the High-Capacity cDNA Reverse Transcription Kit (Applied Biosystems). The cycle threshold (Ct) value was measured on a Bio-Rad CFX96 Real-Time System coupled to a C1000 Thermal Cycler (Bio-Rad) using the iTaq Universal SYBR Green Supermix (Bio-Rad). Normalized relative quantities (NRQs) were obtained using the qBase method (*89*), with RPS26E and PAC1 as reference genes for normalization across samples. NRQ values were normalized to the mean value obtained in wild-type or control (EV) plants. For the synthetic PSY1 time-course experiment, NRQ values were normalized to the mean value obtained in Mock-0hs. NRQ Melting curve analyses at the end of the process and “no template controls” were performed to ensure product-specific amplification without primer– dimer artifacts. Primer sequences are given in Supplemental Table 3. Three biological replicates were analyzed.

### Statistical Analysis

Statistical analysis was performed using GraphPad Prism 9 (GraphPad Software). The specific statistical tests ran are specified in the figure legends. Differences were considered to be significant when P < 0.05.

## Acknowledgments

We thank Jorge Dubcovsky for the use of the Zeiss stereoscope; Ben Scheres for the *ProPLT1-CFP* and *ProPLT1:PLT1-YFP* reporters; Gloria Muday for the *ProCHS:CHS-GFP* reporter; Christoph Ringli for the *ProFLS1:FLS1-GFP* reporter; Wolfgang Busch for the *DR5v2:3nGFP* reporter; Cris Argueso for the p*TCSn::GFP* reporter, Peter Doener for the *ProCYCB1;1:CYCB1;1-GFP* reporter, Audrey Adamchak, Rory Greenhalgh, Ryan Packer and Tracy Weitz for their invaluable help, Ellen Youngsoo Rim, Valley Stewart, Dee Dee Luu, Wolfgang Busch and Ramiro E. Rodriguez for the comments on the manuscript.

## Funding

M.F.E. is a Latin American Fellow in the Biomedical Sciences supported by the Pew Charitable Trusts.

A.M.S. is a USDA-NIFA-AFRI Postdoctoral Fellow (2023-67012-39889).

Innovative Genomics Institute (IGI), funded by the Chan Zuckerberg Initiative National Institutes of Health (GM148173)

Joint Bioenergy Institute, funded by the US Department of Energy (no. DE-AC02-05CH11231). The RNA-seq analysis (proposal: https://doi.org/10.46936/10.25585/60000897) was conducted by the U.S. Department of Energy Joint Genome Institute, a DOE Office of Science User Facility, supported by the Office of Science of the U.S. Department of Energy operated under Contract No. DE-AC02-05CH11231.

National Institutes of Health Administrative Supplements for the purchase of the Confocal Microscope Leica SP8 (Grant no. GM122968).

## Author contributions

Conceptualization: MFE, PCR

Methodology: MFE, AMS, ATAJr, RJ, PCR

Investigation: MFE, AMS, ATAJr, RJ, PCR

Visualization: MFE, PCR

Supervision: PCR

Writing—original draft: MFE, PCR

Writing—review & editing: MFE, AMS, ATAJr, RJ, PCR

## Competing interests

The authors declare that they have no competing interests.

## Data and materials availability

All data are available in the main text or the supplementary materials. The raw data for RNA sequencing are available in the Sequence Read Archive database under BioProject PRJNA616620, PRJNA616621, PRJNA616622, PRJNA616623, PRJNA616551, PRJNA616552 . A previous version of this work was deposited in the preprint depository server bioRxiv. All code from this study is available upon request.

## Supplementary Materials

Figs S1 to S10

Tables S1 to S3

Data S1 to S4

## Notes

### Competing Interest Statement

The authors have declared no competing interest.

