## Supplemental figures and tables for "Tyrosine-sulfated peptide hormone induces flavonol biosynthesis to control elongation and differentiation in Arabidopsis primary root"

Maria Florencia Ercoli *et al.*

**This PDF file includes:**

Figs. S1 to S10
Tables S1 to S3

**Other Supplementary Materials for this manuscript include the following:**
Data S1 to S4

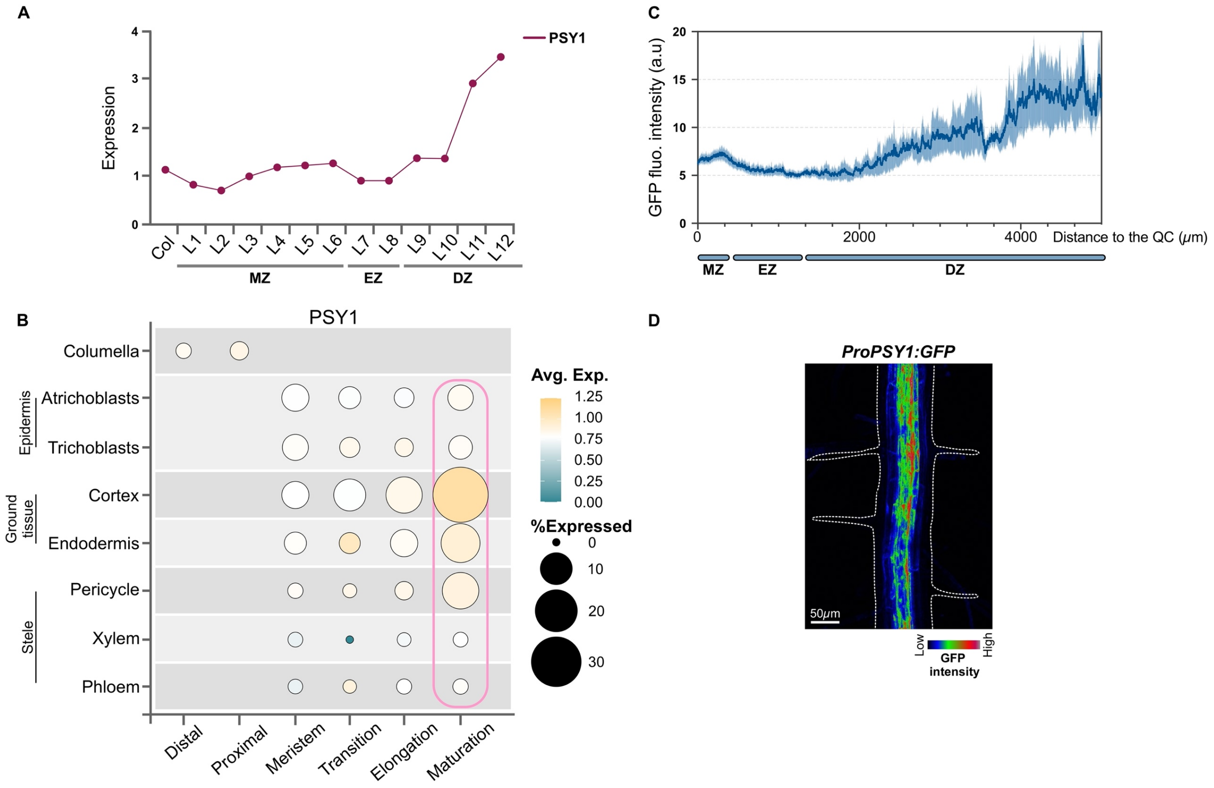


**Supplementary Figure S1. PSY1 expression pattern in Arabidopsis roots.**
Expression profiles of *PSY1* in the root longitudinal axis using data extracted from (**A**) the RootMap database (34) and (**B**) the single-cell Arabidopsis root atlas (35). In (**B**), the developmental stage expression profile for PSY1 is depicted across the four major root tissue types. Dot size represents the percentage of cells at a given developmental stage in which PSY1 is expressed (% Expressed). Dot colors indicate the averaged normalized gene expression value of PSY1 in each developmental stage and cell type group, with warmer colors indicating higher expression levels. The pink square highlights the tissues imaged in (**D**). The developmental stages depicted in (**B**) include Distal and Proximal (only in columella), Meristem (MZ), Transition (TZ), Elongation (EZ) and Maturation (DZ). (**C**) Plot profiles of GFP fluorescence intensity (arbitrary units) in 6-day-old wt seedlings expressing *ProPSY1:GFP*. Data are mean ± s.e.m. of n=10 independent transgenic lines expressing the *ProPSY1:GFP* construct. (**D**) *ProPSY1:GFP* expression in the differentiation zone (4500μm from the QC). This image is a maximum intensity projection generated from 30 z-section images. GFP fluorescence intensity profile is shown using RGB-rainbow false color with its color scale bar, white dashed lines mark organ boundaries. MZ= meristematic zone, EZ=elongation zone, and DZ=differentiation zone.

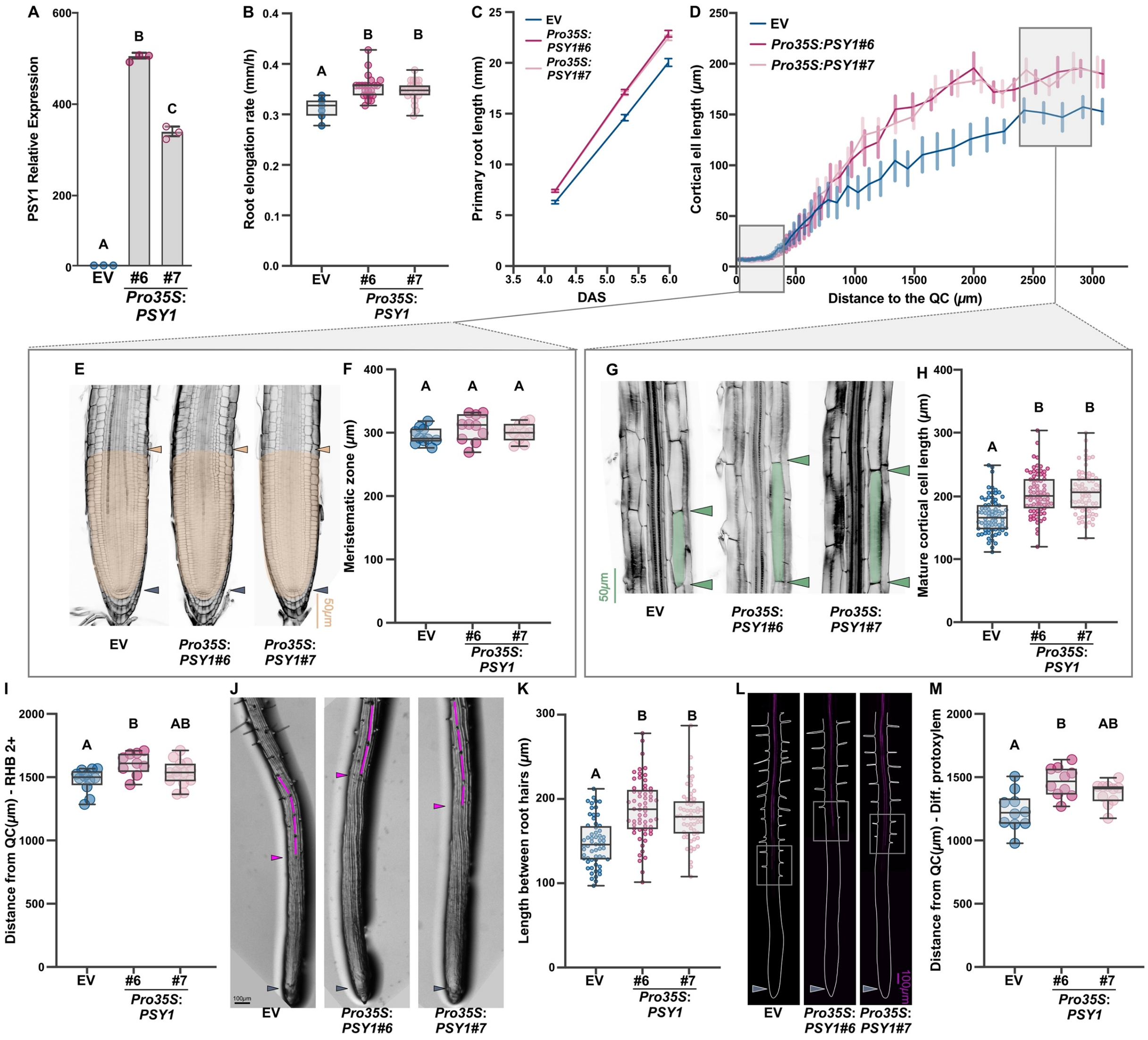


**Supplementary Figure S2. PSY1 acts on cell elongation.**

(**A**) Expression of PSY1 in two independent homozygous transgenic lines *Pro35S:PSY1*#6 and *Pro35S:PSY1*#7 using plants expressing an EV (empty vector) as a control. Expression was estimated by RT-qPCR in three biological replicates and normalized to the mean value obtained in wt plants. (**B**) Root elongation rate (mm/h) (**n=23** seedlings), (**C**) root growth, and (**D**) cortical cell length profile (**n=10** seedlings) in 7-day-old independent homozygous transgenic lines. The meristematic zone size (**E** and **F, n=10** seedlings) and mature cortical cell length (**G** and **H**, **n=75** cells) are highlighted. The meristematic zone (**E**) and mature cortex cells (**F**) are shaded in pale orange and green, respectively. (**I**) Distance from QC to first root hair bulge at stage +2 (RHB 2+) (**n=10-14** seedlings), (**J**) root tip architecture, (**K**) length between consecutive root hairs in one trichoblast file (**n=60**), (**L** and **M**, **n=10** seedlings) distance from QC to differentiated vascular elements (Diff. protoxylem) revealed by basic fuchsin staining in 7-day-old independent homozygous transgenic lines (*Pro35S:PSY1*#6 and #7) with EV (empty vector) control. In (**J**), the length between consecutive root hairs in one trichoblast file is highlighted in magenta. In (**l**), the grey squares indicate the zone where the deposition of lignin starts in the protoxylem, as revealed with basic fuchsin staining, and organ boundaries are marked by white dashed lines. In (**B**), (**F**), (**H**), (**I**), (**K**), and (**M**), the data shown are box and whisker plots combined with scatter plots; each dot indicates the measurement of the designated parameter listed on the y-axis of the plot. Different letters indicate significant differences, as determined by one-way ANOVA followed by Tukey's multiple comparison test ($P < 0.05$). The purple arrowheads mark the position of the QC, the pale orange arrowheads mark the end of the meristem, where cells start to elongate, the green arrowheads indicate the mature cortical cell size, and the magenta arrowhead points to the first root hair bulge, defined as stage +2, indicating the end of the elongation zone in the epidermis.

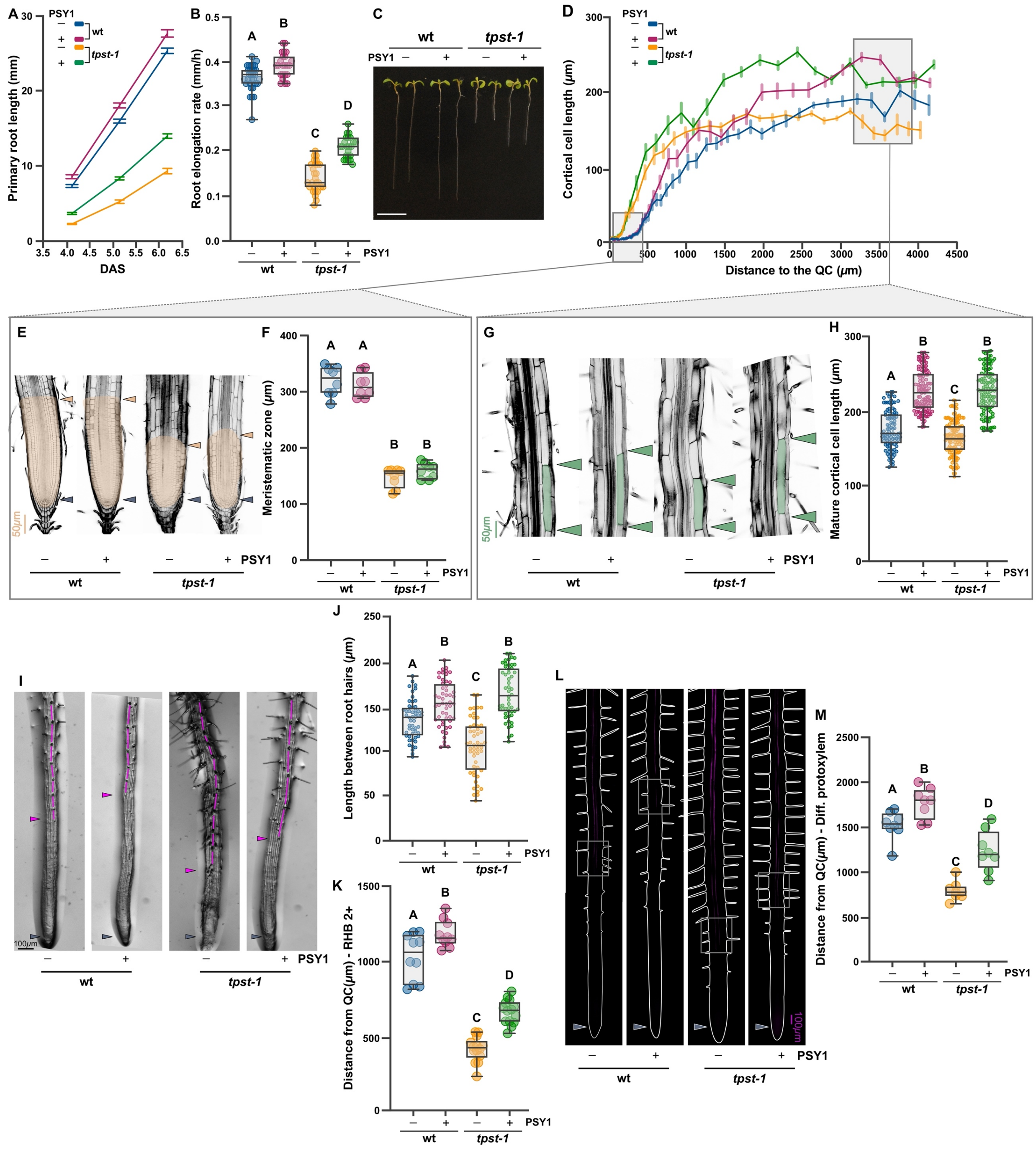


**Supplementary Figure S3. Synthetic PSY1 treatment rescues the cell elongation defects in *tpst-1* plants.**

(**A**) Root growth, (**B**) root elongation rate (mm/h)(**n=23** seedlings), (**C**) root phenotype, and (**D**) cortical cell length profile (**n=8** seedlings) in wt and *tpst-1* seedlings grown for 6 days on 1X MS vertical plates with or without 50nM of PSY1. The meristematic zone size (**E** and **F, n=8** seedlings) and mature cortical cell length (**G** and **H**, **n=100** cells) are highlighted. The meristematic zone (**E**) and mature cortex cells (**G**) are shaded in pale orange and green, respectively. (**I**) Root tip architecture, (**J**) length between consecutive root hairs in one trichoblast file (**n=52**), (**K**) distance from QC to first root hair bulge at stage +2 (RHB 2+) (**n=10-14** seedlings), (**L** and **M, n=8** seedlings) distance from QC to differentiated vascular elements (Diff. protoxylem) revealed by basic fuchsin staining in wt and *tpst-1* seedlings grown for 6 days on 1X MS vertical plates with or without 50nM of PSY1. In (**I**), the length between consecutive root hairs in one trichoblast file is highlighted in magenta. In (**L**), the grey squares indicate the zone where the deposition of lignin starts in the protoxylem, as revealed with basic fuchsin staining, and organ boundaries are marked by white dashed lines. In (**B**), (**F**), (**H**), (**J**), (**K**), and (**M**), the data shown are box and whisker plots combined with scatter plots; each dot indicates the measurement of the designated parameter listed on the y-axis of the plot. Different letters indicate significant differences, as determined by 2-way ANOVA followed by Tukey's multiple comparison test ($P < 0.05$). The purple arrowheads mark the position of the QC, the pale orange arrowheads mark the end of the meristem, where cells start to elongate, the green arrowheads indicate the mature cortical cell size, and the magenta arrowhead points to the first root hair bulge, defined as stage +2, indicating the end of the elongation zone in the epidermis.

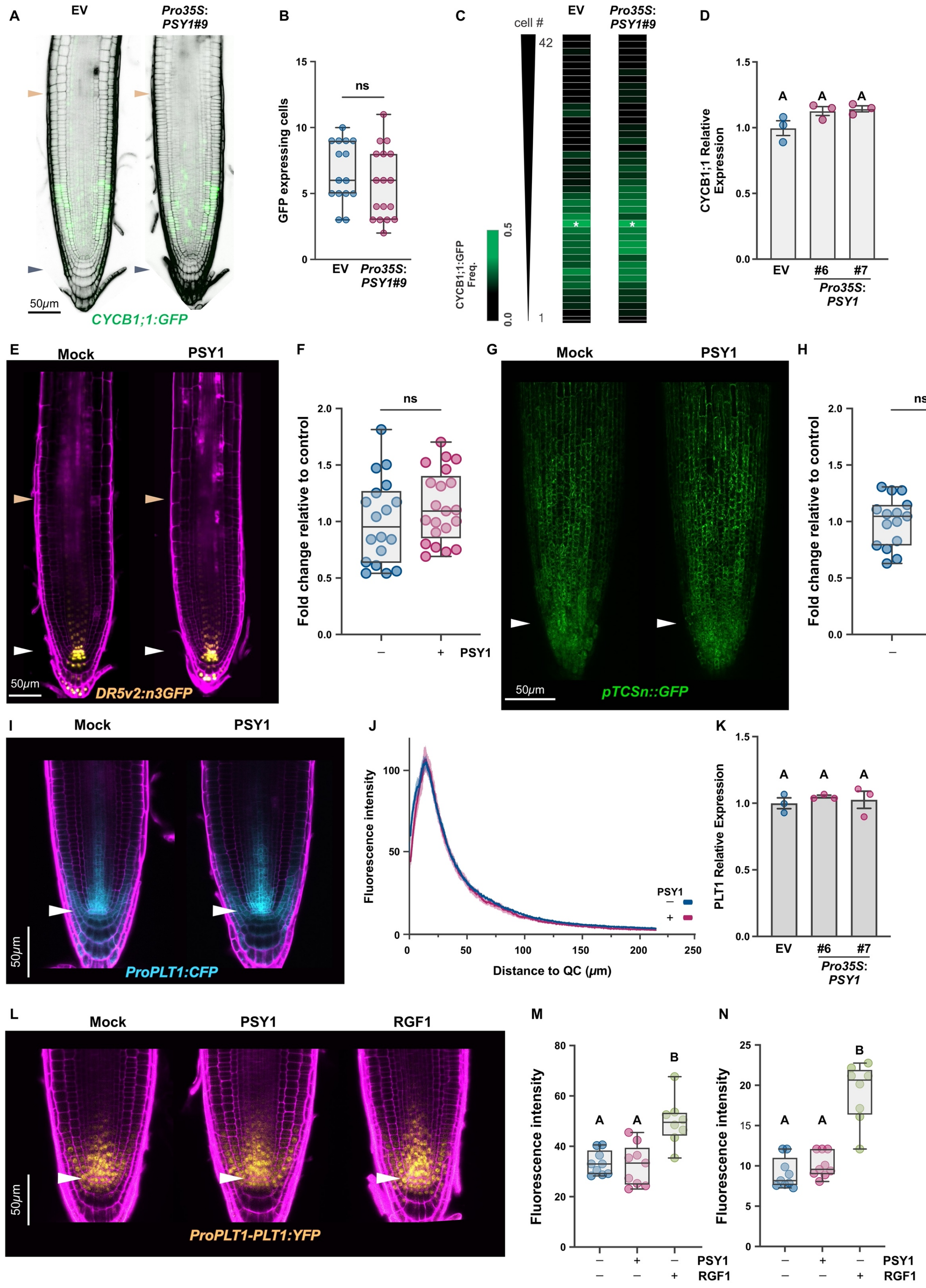


**Supplementary Figure S4. Cell proliferation in Arabidopsis roots is not altered by PSY1.** (**A**) Localization of dividing cells in root apical meristems and (**B**) GFP expressing cells per cortex cell files in a *CYCLINB1;1* reporter line *(CYCB1;1:GFP*) transformed with *Pro35S:PSY1* or EV (empty vector) as control. *Pro35S:PSY1#9* is a homozygous transgenic line that was selected as a representative of several independent transgenic lines analyzed. (**C**) Heat map showing the frequency of *CYCB1;1:GFP*-positive cells at a given distance from the QC quantified by the number of cortical cells. Note that the maximum frequencies (star) occur at the same distance from the QC in EV and *Pro35S:PSY1#9* roots. The distribution of *CYCB1;1: GFP*-expressing cells was scored from the cell adjacent to the QC (1) up to cell 40. Thirty-six cortex cell files from 18 seedlings for each genotype were scored. (**D**) Expression of *CYCB1;1* in two independent homozygous transgenic lines *Pro35S:PSY1*#6 and *Pro35S:PSY1*#7 using EV (empty vector) as control. Expression was estimated by RT-qPCR in three biological replicates and normalized to the mean value obtained in wild-type (wt) plants. Expression pattern and fluorescence intensity quantification of the auxin response reporter line *DR5v2:n3GFP* (yellow) (**E, F**) and cytokinin response reporter line *pTCSn::GFP* (green) (**G, H**) in the meristematic zone of seedlings grown for 6 days on 1xMS vertical plates with or without 100nM of synthetic PSY1. In (**F**) and (**H**), the fluorescence intensity is plotted as a fold change relative to the control. In (**B**),(**F**), and (**H**), the data shown are box and whisker plots combined with scatter plots; each dot indicates the measurement of the designated parameter in an independent seedling, ns indicates no significant differences (Student's t-test, $p>0.05$). (**I**) Expression of the reporter line *ProPLT1:CFP* (blue), and (**J**) its fluorescence intensity profile in root tips in wt seedlings grown for 6 days on 1xMS vertical plates with or without 100nM of synthetic PSY1. Fluorescence intensity is expressed in arbitrary units. (**K**) Expression of *PLT1* in two independent homozygous transgenic lines *Pro35S:PSY1*#6 and *Pro35S:PSY1*#7 using EV (empty vector) as control. Expression was estimated by RT-qPCR in three biological replicates and normalized to the mean value obtained in wt plants. (**L**) *ProPLT1-PLT1:YFP* (yellow) expression gradient is modified by RGF1 but not by PSY1 treatment. (**M**) and (**N**) are quantifications of the fluorescence intensity of proteins produced by *ProPLT1-PLT1:YFP* reporter in wt seedlings grown for 6 days on 1xMS vertical plates with or without 100nM of synthetic PSY1. For RGF1 treatments, 5-day-old seedlings were transferred to 1xMS vertical plates with or without 100nM of synthetic RGF1 for 24 hours. In (**M**), measurements were done at the QC region, while in (**N**), the stele region 50 um above QC was measured. In (**M**) and (**N**), the data shown are box and whisker plots combined with scatter plots; each dot indicates the measurement of the designated parameter in an independent seedling, and fluorescence intensity is expressed in arbitrary units. Different letters indicate significant differences, as determined by ANOVA followed by Tukey's multiple comparison test ($P < 0.05$). The purple and white arrowheads mark the position of the QC, and the pale orange arrowheads mark the end of the meristem, where cells start to elongate.

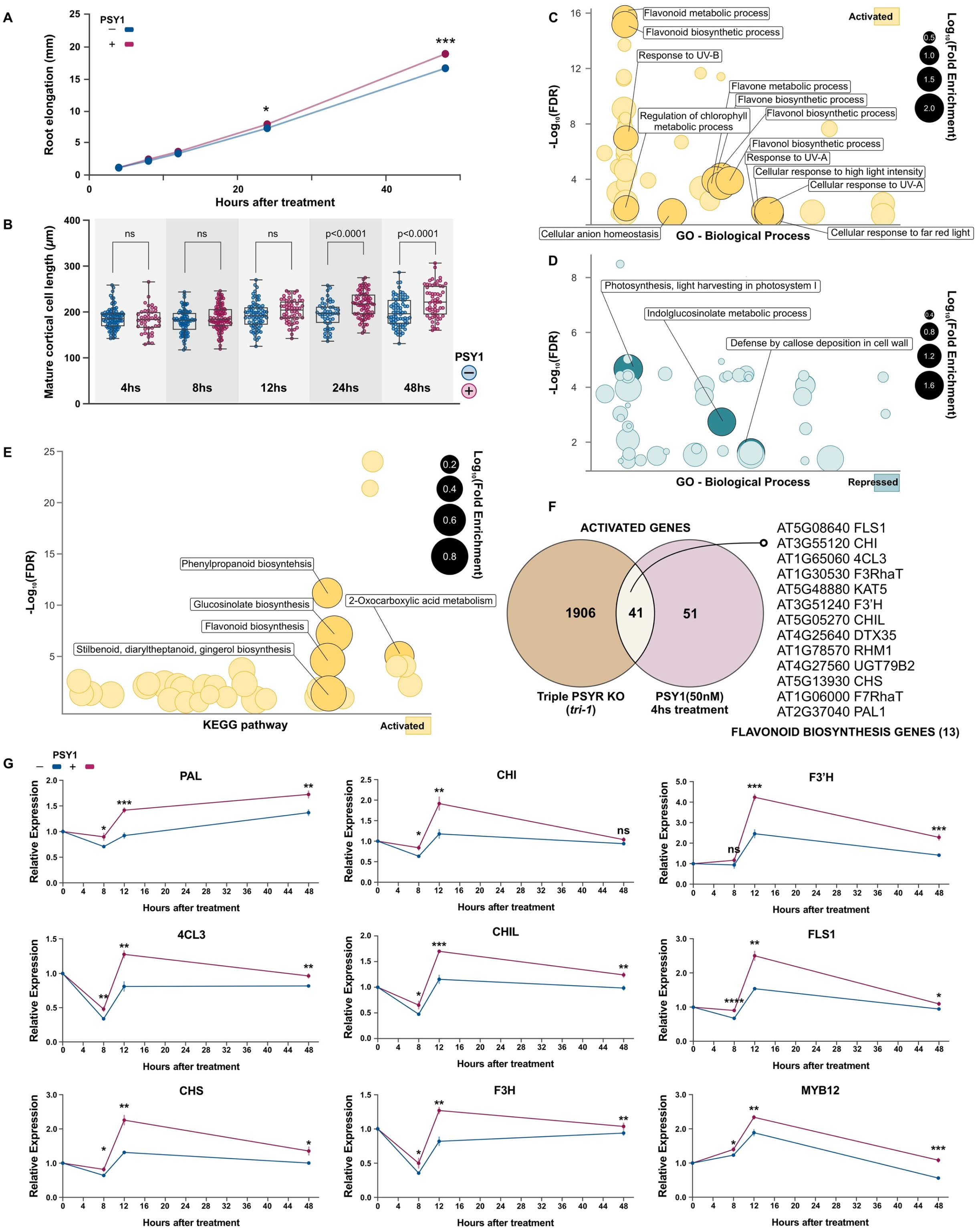


**Supplementary Figure S5. Flavonoid biosynthesis genes are activated by PSY1.**
(**A**) Root growth and (**B**) mature cortical cell length in wt 7-day-old seedlings treated with PSY1 for 4, 8, 12, 24, and 48 hours. Bubble plots for GO-Biological Process significant terms for genes

(**C**) activated and (**D**) repressed by PSY1 (Supplemental Data set 2). (**E**) Bubble plot for Kyoto Encyclopedia of Genes and Genomes (KEGG) pathway enrichment analysis of genes activated in the triple PSYR mutant (*tri-1*) (Supplemental Data set 4). In (**C**),(**D**), and (**E**), the y-axis shows the False Discovery Rate (FDR) in a negative Log10 scale, whereas the x-axis is fixed, and terms from the same GO/KEGG subtree are located closer to each other. The size of each circle represents the term Fold Enrichment in the Log10 scale. In (**C**) and (**D**), GO terms with a Log10 (Fold Enrichment) higher than 1.5 are highlighted. In (**E**), KEGG pathways with a Log10 (Fold Enrichment) higher than 0.5 are highlighted. (**F**) Venn diagram depicting the overlap between activated genes in the triple PSYR mutant (*tri-1*) from (Wang et al., 2022) and PSY1 4-hour treatment. 13 of the 41 genes in the overlap are involved in flavonoid biosynthesis. (**G**) Expression of selected PSY1 target genes in wt 7-day-old seedlings upon treatment with 250nM PSY1 for 8,12, and 48 hours, measured by quantitative RT–PCR. The samples were generated using complete seedlings, including shoots. Expression was estimated by RT-qPCR in three biological replicates and normalized to the mean value obtained in seedlings grown in the absence of PSY1 at time 0 hours. Data shown are means +/- SE of three biological replicates. P values are calculated by two-tailed Student's t-test (*$P \leq 0.05$, **$P \leq 0.01$, *** $P \leq 0.001$, **** $P \leq 0.0001$). The selected activated genes included *PAL,* phenylalanine ammonia-lyase*; 4CL3,* 4-coumaric acid: CoA ligase 3; *CHS,* chalcone synthase; *CHI,* chalcone isomerase; *CHIL,* chalcone isomerase-like, *F3H,* flavanone 3-hydroxylase; *F3'H,* flavonoid 3-hydroxylase; *FLS1*, flavonol synthase*,* and the transcription factor *MYB12.*

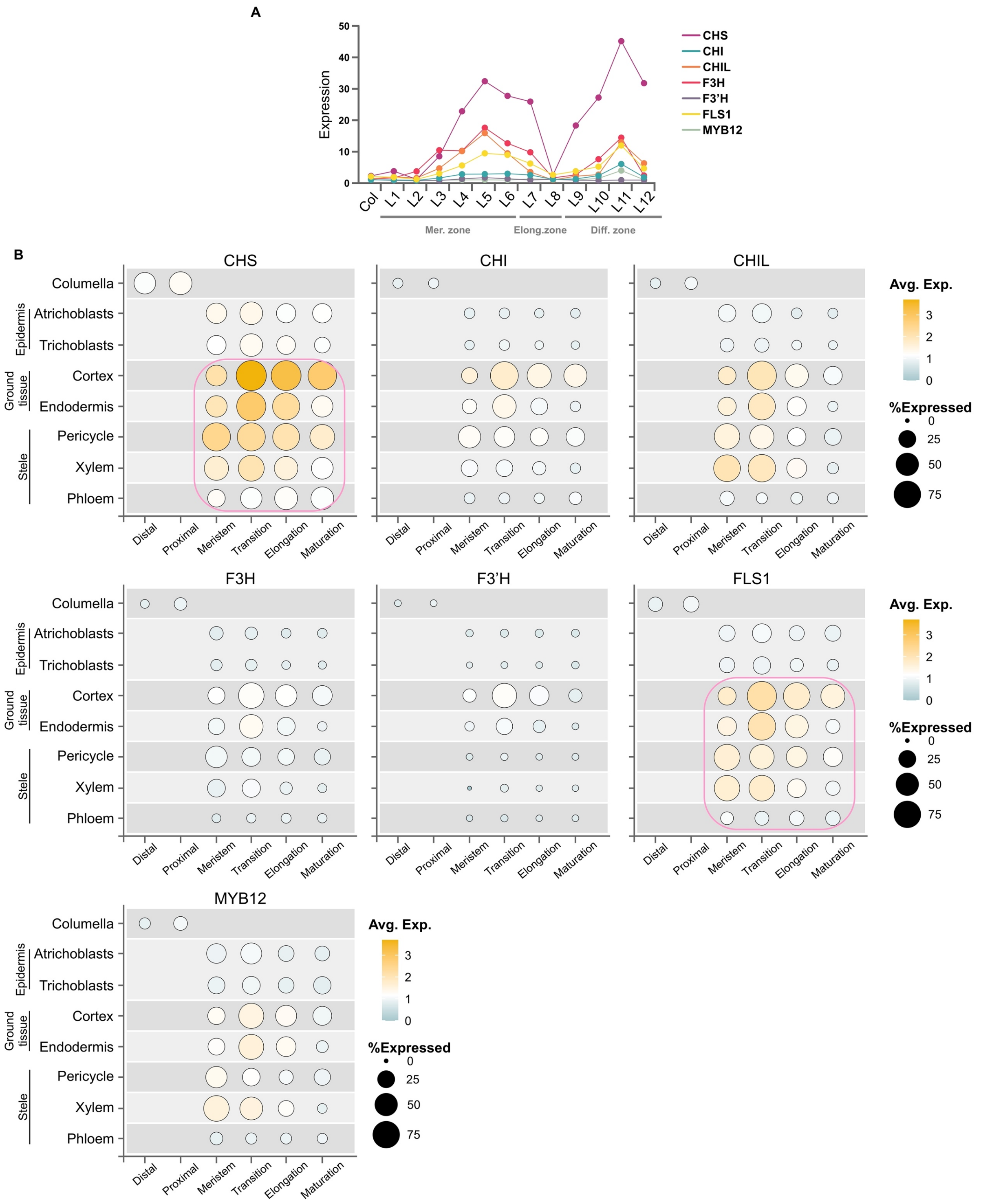


**Supplementary Figure S6. Expression pattern along the Arabidopsis primary root of early flavonoid biosynthesis genes.**

Expression profiles of 7 genes that participate in the flavonoid in the root longitudinal axis using data extracted (**A**) the RootMap database (34) and (**B**) the single-cell Arabidopsis root atlas (35).

In **(B)**, the developmental stage expression profile for the selected genes is depicted across the four major root tissue types. Dot size represents the percentage of cells at a given developmental stage in which each gene is expressed (% Expressed). Dot colors indicate the averaged normalized gene expression value of each gene in each developmental stage and cell type group, with warmer colors indicating higher expression levels. The developmental stages depicted in (**B**) include Distal and Proximal (only in columella), Meristem (MZ), Transition (TZ), Elongation (EZ) and Maturation (DZ). The pink square in CHS and FLS1 highlights the tissues for which high expression has been confirmed using reporter lines (52). *CHS,* chalcone synthase; *CHI,* chalcone isomerase; *CHIL,* chalcone isomerase-like, *F3H,* flavanone 3-hydroxylase; *F3'H,* flavonoid 3-hydroxylase; *FLS1*, flavonol synthase1*,* and the transcription factor *MYB12.*

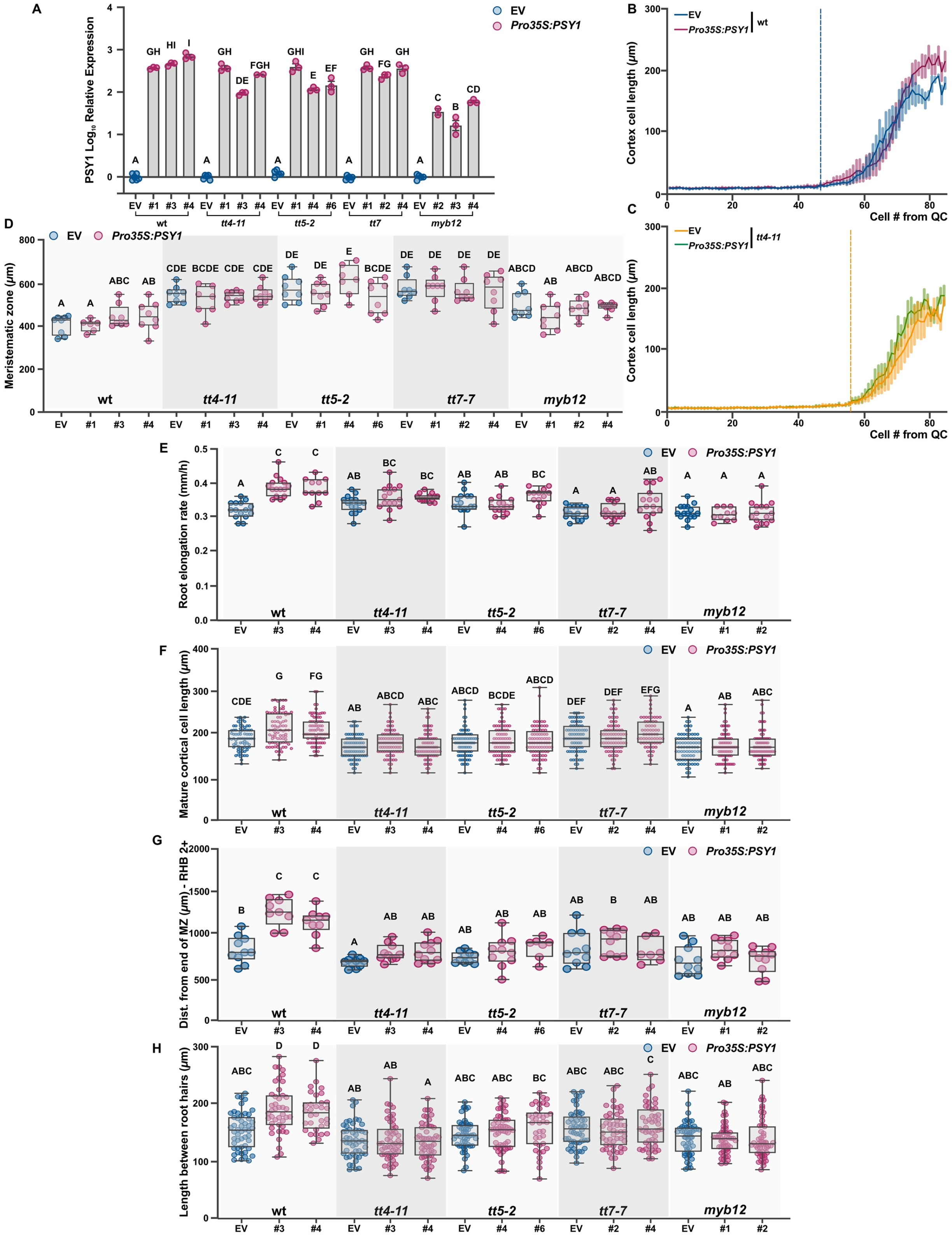


**Supplementary Figure S7. Mutants in the flavonoid biosynthetic pathway are less responsive to PSY1 accumulation.**

(**A**) Expression of PSY1 (Log10 scale) in three independent homozygous transgenic lines (*Pro35S: PSY1*) with EV (empty vector) control generated in wild-type (wt-Col0) and mutants defective in flavonol biosynthesis (*tt4-11, tt5-2,tt7-7 and myb12*) backgrounds. Expression was estimated by RT-qPCR in three biological replicates and normalized to the mean value obtained EV control plants for each background. Different letters indicate significant differences, as determined by one-way ANOVA followed by Tukey's multiple comparison test (P < 0.05). (**B** and **C**) cortical cell length profile (**n=8** seedlings) in 7-day-old independent homozygous transgenic lines that accumulated higher levels of PSY1 with EV (empty vector) control generated in wt (**B**) and *tt4-11* (**C**). The dotted line indicates the end of the meristematic zone, highlighting the longer meristematic zone in *tt4-11* compared to the control. (**D**) Meristematic zone size (**n=8** seedlings), (**E**) Root elongation rate (mm/h)(**n=10-15** seedlings), (**F**) mature cortical cell length (**n=80** cells), (**G**) distance from the end of the MZ to first root hair bulge at stage +2 (RHB 2+) (**n=6-10** seedlings), (**H**) length between consecutive root hairs in one trichoblast file (**n=50**) in 7-day-old independent homozygous transgenic lines that accumulated higher levels of PSY1 (*Pro35S: PSY1*) with EV (empty vector) control generated in wt (wt-Col-0) and mutants defective in flavonol biosynthesis (*tt4-11, tt5-2,tt7-7 and myb12*) backgrounds. In (**D**), (**E**),(**F**),(**G**), and (**H**), the data shown are box and whisker plots combined with scatter plots; each dot indicates the measurement of the designated parameter listed on the y-axis of the plot. Different letters indicate significant differences, as determined by one-way ANOVA followed by Tukey's multiple comparison test (P < 0.05). Different numbers indicate independent transgenic lines overexpressing PSY1.

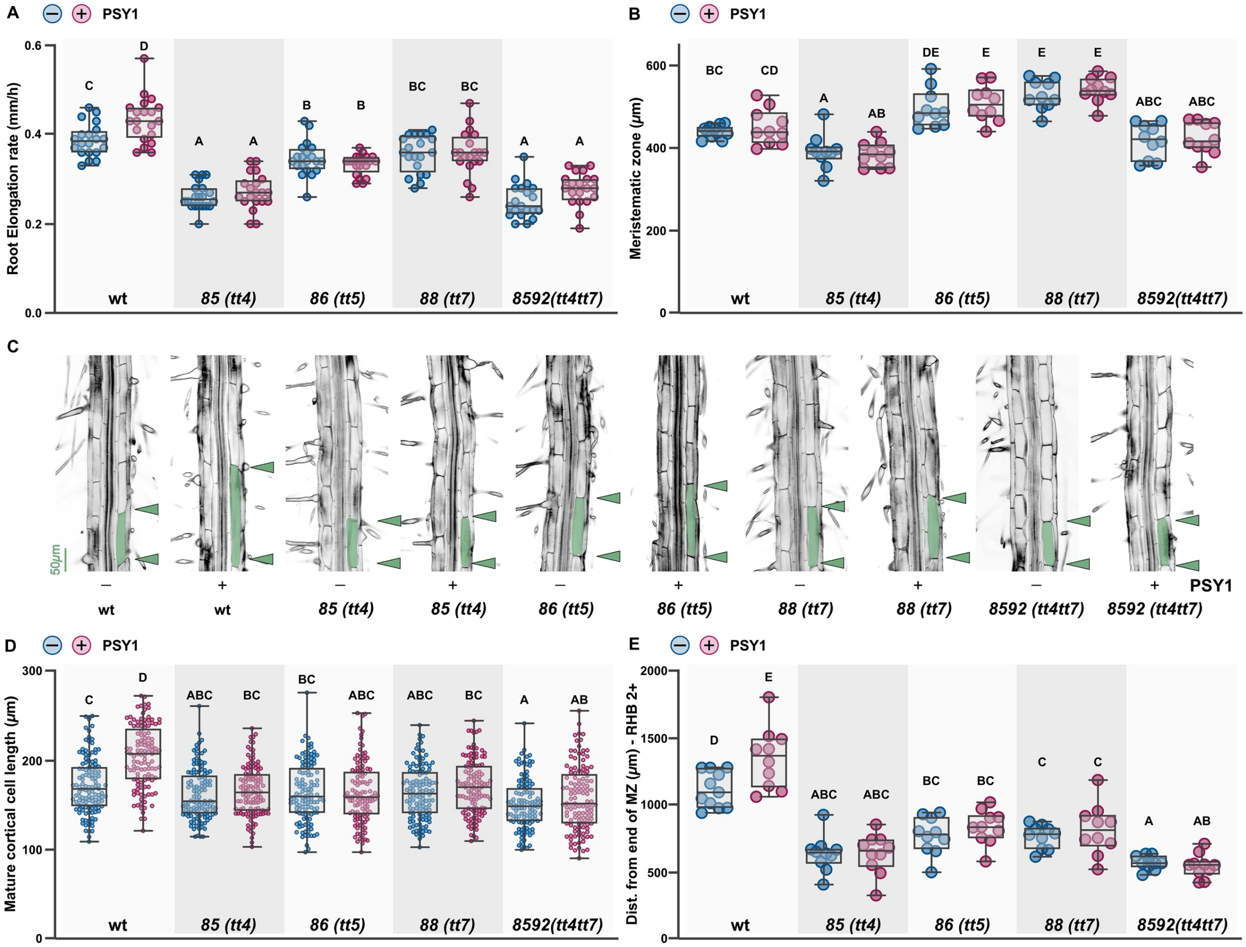


**Supplementary Figure S8. PSY1-induced root growth is absent in flavonoid biosynthetic mutants available in a different accession.**

(**A**) Root elongation rate (mm/h) (**n=20** seedlings), (**B**) Meristematic zone size (**n=10** seedlings), (**C**) distance from QC to first root hair bulge at stage +2 (RHB 2+) (**n=10** seedlings), (**D** and **E**) mature cortical cell length (**n=120** cells), in wild-type (wt-Ler) and mutants defective in flavonol biosynthesis (*85 or tt4, 86 or tt5, 88 or tt7 and 8592 or tt4tt7*) seedlings grown for 6 days on 1xMS vertical plates with or without 100nM of PSY1. In (**D**), the limits of representative mature cortex cells are shaded in green with green arrowheads indicating the mature cortical cell size. In (**A**), (**B**),(**C**), and (**E**), the data shown are box and whisker plots combined with scatter plots; each dot indicates the measurement of the designated parameter listed on the y-axis of the plot. Different letters indicate significant differences, as determined by one-way ANOVA followed by Tukey's multiple comparison test ($P < 0.05$).

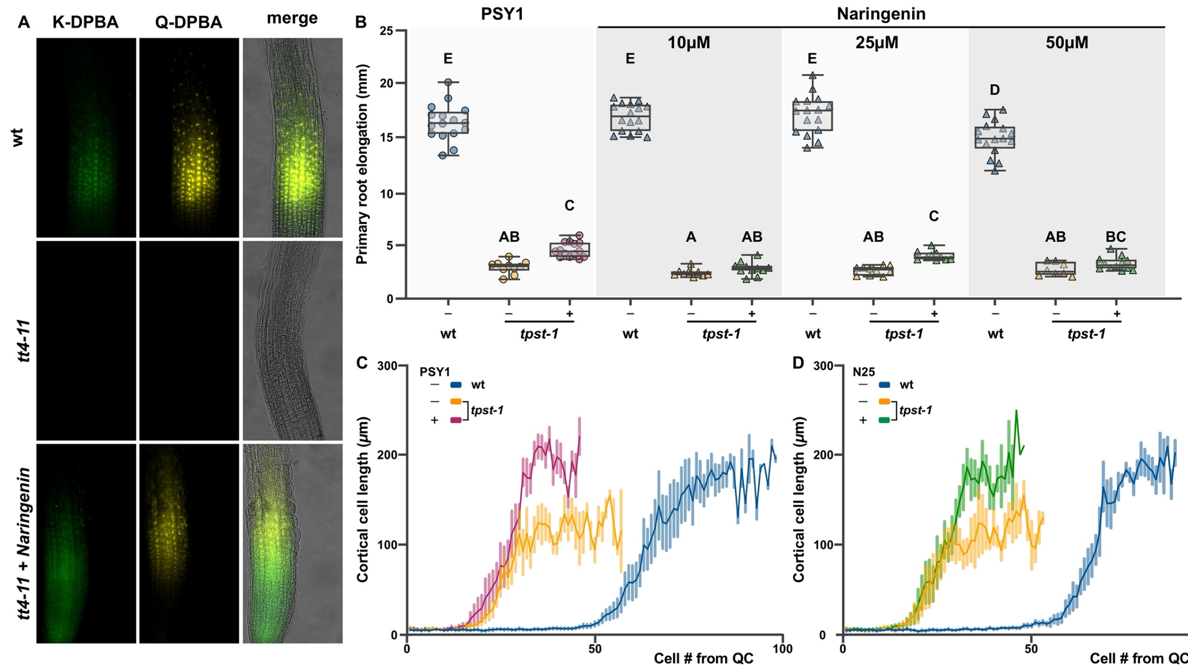


**Supplementary Figure S9. Naringenin synthetic treatment induces root growth in *tpst-1*.**
(**A**) Confocal images of 5-day-old DPBA-stained wild-type (wt, Col0) and *tt4-11* primary roots before and 6 hours after treatment with 25µM Naringenin (see Figure 2 for biosynthetic pathway). (**B**) Root elongation (**n=10-16** seedlings) in 5-day-old wt and *tpst-1* seedlings grown for 48hs on control conditions (1X MS or 1X MS supplemented with EtOH) or treated with 50nM PSY or 10,25, or 50µM Naringenin. The data is shown as box and whisker plots combined with scatter plots; each dot indicates the individual measurements of root elongation. Different letters indicate significant differences, as determined by one-way ANOVA followed by Tukey's multiple comparison test ($P < 0.05$). (**C**) Cortical cell length profile (**n=8** seedlings) in 5-day-old wt and *tpst-1* seedlings grown for 48hs on control conditions (1X MS or 1X MS supplemented with EtOH) or treated with 50nM PSY or 25µM Naringenin.

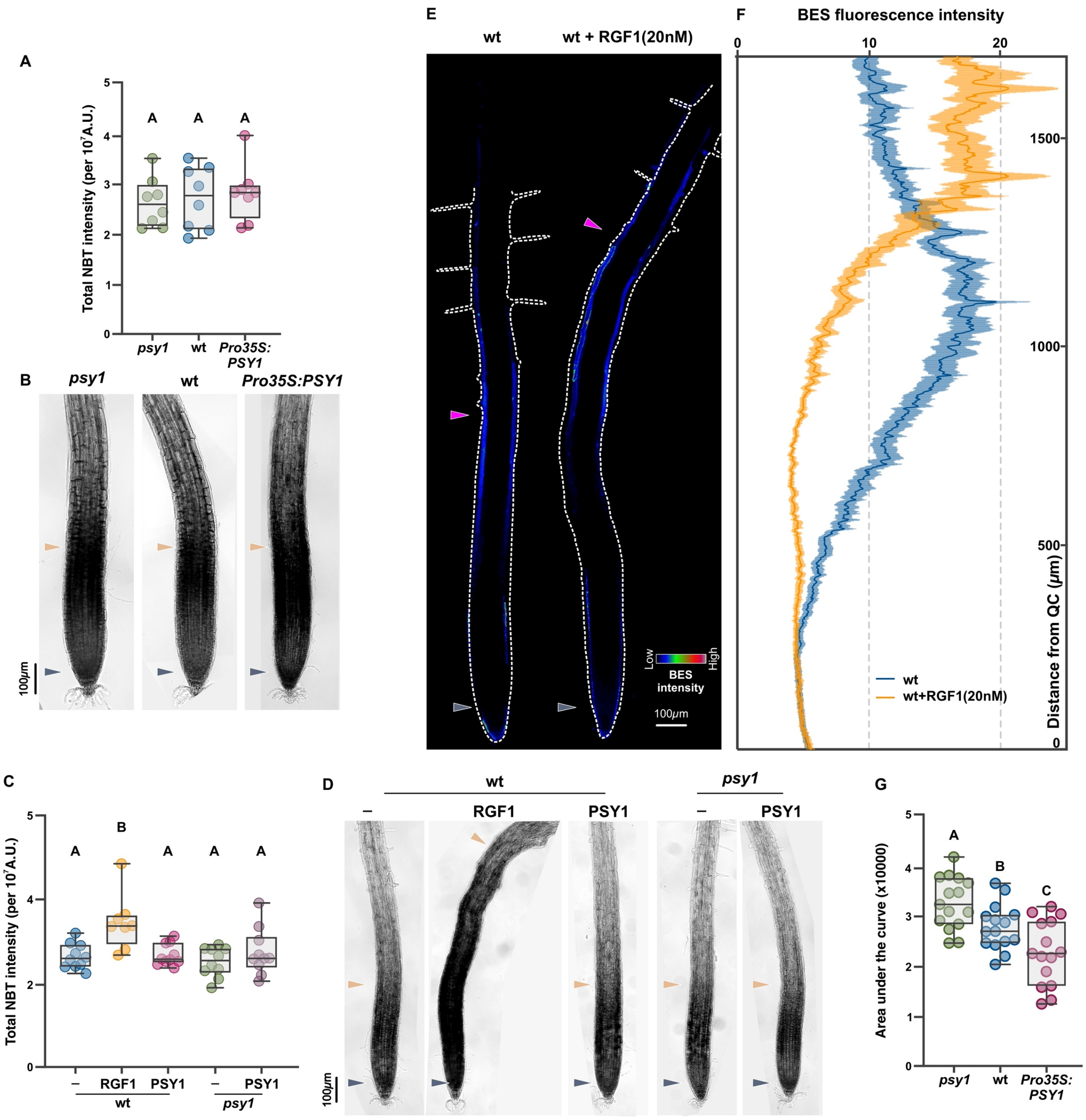

**Supplementary Figure S10. PSY1 does not affect $O_2^-$ accumulation in the meristematic zone.**

(**A**) and (**C**) shows quantification of NBT staining intensity (arbitrary units) in the meristematic zone from (**B**) and (**D**), respectively (**n= 8-10** seedlings). (**B**) NBT staining of 7-day-old *psy1*, wt, and *Pro35S:PSY1* seedlings. (**D**) NBT staining of 5-day-old wt and *psy1* seedlings after 48-hour treatment with or without RGF1 20nM or PSY1 100nM. (**E**) Representative epidermal BES fluorescence 5-day-old wt seedlings after 24hs treatment with or without RGF1 20nM. BES fluorescence intensity profile is shown using RGB-rainbow false color with its color scale bar; white dashed lines mark organ boundaries. Images are scaled to correspond to the y-axis of the plot profile in (**F**). (**F**) Plot profiles of epidermal BES fluorescence in 5-day-old wt seedlings after 24-hour treatment with RGF1 20nM. Data are mean ± s.e.m. of **n=8** seedlings. (**G**) The area under the curve from plot profiles of epidermal BES fluorescence in *psy1*, wt, and *Pro35S:PSY1*

6-day-old seedlings (**n=15** seedlings). In (**A**), (**C**), and (**G**), the data shown are box and whisker plots combined with scatter plots; each dot indicates the measurement of the designated parameter. Different letters indicate significant differences, as determined by one-way ANOVA followed by Tukey's multiple comparison test ($P < 0.05$). The purple arrowheads mark the position of the QC, the pale orange arrowheads mark the end of the meristem, where cells start to elongate, and the magenta arrowhead points to the first root hair bulge, defined as stage +2, indicating the end of the elongation zone in the epidermis.

**Table S1. Arabidopsis mutant lines used in this study.**

| Gene product, locus, mutant name | Allele | Ecotype<br>Mutation type | First characterized in |
|---|---|---|---|
| **PLANT PEPTIDE SULFATED IN TYROSINE 1** (PSY1)<br>AT5G58650<br>*psy1* | *psy1-1* | Col0<br>GK-583C09 | (*33*) |
| **TYROSYLPROTEIN SULFOTRANSFERASE** (TPST)<br>AT1G08030<br>*tpst* | *tpst-1* | Col0<br>SALK_009847 | (*25*) |
| **CHALCONE SYNTHASE** (CHS)<br>AT5G13930<br>*tt4* | *tt4-11* | Col0<br>SALK_020583 | (*90*) |
| | *85 (tt4)*<br>*8592(tt4tt7)* | Ler<br>EMS | (*91*) |
| **CHALCONE ISOMERASE** (CHI)<br>AT3G55120<br>*tt5* | *tt5-2* | Col0<br>GK-176H03 | (*92*) |
| | *86(tt5)* | Ler<br>Fast neutrons | (*93*) |
| **FLAVONOID 3′ HYDROXYLASE** (F3′H)<br>AT5G07990<br>*tt7* | *tt7-7* | Col0<br>GK-629C11 | (*54*) |
| | *88 (tt7)*<br>*8592(tt4tt7)* | Ler<br>EMS | (*94*) |
| **PRODUCTION OF FLAVONOL GLYCOSIDES 1** AT2G47460<br>*myb12* | *myb12* | Col0<br>SALK_046675C | (*95*) |

**Table S2. Arabidopsis reporter lines used in this study.**

| Reporter line | Description | Reference |
|---|---|---|
| *CYCB1;1:GFP* | CYCLINB1;1 reporter line (G2-M specific marker) | (*96*) |
| *ProPLT1:CFP* | PLT1 transcriptional reporter line | (*16*) |
| *ProPLT1:PLT1-YPF* | PLT1 translational reporter line | (*16*) |
| *DR5v2:3nGFP* | Synthetic auxin-inducible promoter driving the expression of nuclear-localized GFP | (*38*) |
| *pTCSn::GFP* | Two Component signaling Sensor (TCS)::green fluorescent protein (GFP), which reflects the transcriptional activity of type-B response regulators | (*97*) |
| *ProCHS:CHS-GFP* | CHS translational reporter line | (*53*) |
| *ProFLS1:GFP-FLS1* | FLS1 translational reporter line | (*98*) |

**Table S3. Oligonucleotide primers used in this study.**

| Gene | Locus ID | Sequence (5' - 3') | Purpose |
|---|---|---|---|
| **PSY1** | AT5G58650 | CACCATGACTTTTGTAGTTCGTC | PSY1 Topo Cloning and genotyping Fw |
| | | AAACATTACGTCAGCCTCTGC | PSY1 Topo Cloning Rv |
| | | CAACCCTGTTTCCGTTTCAGGTG | RT-qPCR primer Fw |
| | | CACCGTAGTCCTCAACGTTCACCA | RT-qPCR primer Rv |
| | | GAAACGCTGGAACCTTTCCGTGAA | *psy1* Genotyping Rv |
| | | ATATTGACCATCATACTCATTGC | o8409_Gabi_TDNA1_LB |
| | | ATAATAACGCTGCGGACATCTACATTTT | o8474_Gabi_TDNA2_LB |
| **ProPSY1** | AT5G58650 | CACCTACGTTCCAAAAAATGA | *Pro:PSY1* Topo Cloning Fw |
| | | ATCTCTCTGTCTGATATAATTAAAAG | *Pro:PSY1* Topo Cloning Rv |
| **PAL1** | AT2G37040 | GACAAGTGGCTGCGATCTCAAC | RT-qPCR primer Fw |
| | | TGCTTCCGAATATTCCGGCGTTAAG | RT-qPCR primer Rv |
| **4CL3** | AT1G65060 | ATGATCACTGCAGCTCTACACGA | RT-qPCR primer Fw |
| | | GAGGCGTAGGAGGAGAATGAGG | RT-qPCR primer Rv |
| **CHS** | AT5G13930 | CCGCATCACCAACAGTGAACAC | RT-qPCR primer Fw |
| | | TCCTCCGTCAGATGCATGTGAC | RT-qPCR primer Rv |
| **CHI** | AT3G55120 | CGGTTCATCGATCCTCTTCGCTC | RT-qPCR primer Fw |
| | | CACAGCGATCCCGGTTTCAGG | RT-qPCR primer Rv |
| **CHIL** | AT5G05270 | CAGCCAAAGATGGCATTTGCGAG | RT-qPCR primer Fw |
| | | ACCATCTCTGCATCATTCCCACC | RT-qPCR primer Rv |
| **F3H** | AT3G51240 | ACGGTTCAGCCTGTTGAAGGAG | RT-qPCR primer Fw |
| | | CCACGGCCTGATGATCAGCA | RT-qPCR primer Rv |
| **F3′H** | AT5G07990 | GGAACTGATCTTGACGGTGACGG | RT-qPCR primer Fw |
| | | GCTGACGTGTCAGTTCCAGCTG | RT-qPCR primer Rv |
| **FLS1** | AT5G08640 | GGAGGTCGAAAGAGTCCAAGACAT | RT-qPCR primer Fw |
| | | GCGTTGGACCTCGGAATGTT | RT-qPCR primer Rv |

| | | | |
|---|---|---|---|
| **MYB12** | AT2G47460 | GAACCAAGGGAATCTCGACTGTCT | RT-qPCR primer Fw |
| | | ACCTTCTTGCCAAACACAATCCCA | RT-qPCR primer Rv |
| **RPS26E** | AT3G56340 | GACTTTCAAGCGCAGGAATGGTG | RT-qPCR primer Fw |
| | | CCTTGTCCTTGGGGCAACACTTT | RT-qPCR primer Rv |
| **PAC1** | AT3G22110 | TCTCTTTGCAGGATGGGACAAGC | RT-qPCR primer Fw |
| | | AGACTGAGCCGCCTGATTGTTTG | RT-qPCR primer Rv |

**Data S1-S4. (separate file)**
